# Phage and bacteria diversification through a prophage acquisition ratchet

**DOI:** 10.1101/2020.04.08.028340

**Authors:** Maxwell Anthenelli, Emily Jasien, Robert Edwards, Barbara Bailey, Ben Felts, Parag Katira, James Nulton, Peter Salamon, Forest Rohwer, Cynthia B. Silveira, Antoni Luque

**Affiliations:** Department of Physics, San Diego State University, San Diego, USA; Department of Mathematics & Statistics, San Diego State University, San Diego, USA; Department of Biology, San Diego State University, San Diego, USA; Department of Mechanical Engineering, San Diego State University, San Diego, USA; Department of Computer Science, San Diego State University, San Diego, USA; Viral Information Institute, San Diego State University, San Diego, USA; Computational Science Research Center, San Diego State University, San Diego, USA

**Author notes:** Co-first authors (Maxwell Anthenelli, Emily Jasien).

## Abstract

Lysogeny is prevalent in the microbial-dense mammalian gut. This contrasts the classical view of lysogeny as a refuge used by phages under poor host growth conditions. Here we hypothesize that as carrying capacity increases, lysogens escape phage top-down control through superinfection exclusion, overcoming the canonical trade-off between competition and resistance. This hypothesis was tested by developing an ecological model that combined lytic and lysogenic communities and a diversification model that estimated the accumulation of prophages in bacterial genomes. The ecological model sampled phage-bacteria traits stochastically for communities ranging from 1 to 1000 phage-bacteria pairs, and it included a fraction of escaping lysogens proportional to the increase in carrying capacity. The diversification model introduced new prophages at each diversification step and estimated the distribution of prophages per bacteria using combinatorics. The ecological model recovered the range of abundances and sublinear relationship between phage and bacteria observed across eleven ecosystems. The diversification model predicted an increase in the number of prophages per genome as bacterial abundances increased, in agreement with the distribution of prophages on 833 genomes from marine and human-associated bacteria. The study of lysogeny presented here offers a framework to interpret viral and microbial abundances and reconciles the Kill-the-Winner and Piggyback-the-Winner paradigms in viral ecology.

## Introduction

The human gut contains one of the highest concentrations of bacteria and phages—viruses that infect bacteria—across ecosystems (Knowles et al. 2016; Wigington et al. 2016; Parikka et al. 2017). This high concentration of microbes is sustained by the daily supply of nutrient-rich compounds received from food intake and microbial metabolism (Blaut 2011; Cotillard et al. 2013; Mirzaei and Maurice 2017). In other ecosystems—mostly aquatic environments—an increase of resources has been linked to bacterial growth and phage lytic life cycle. This phage strategy produces new phage particles upon infection and subsequently bursts the bacterial host (lysis). Combined with the bacterial growth, the lytic life cycle ensures a rapid turnover of nutrients charateristic of the kill-the-winner (KtW) dynamics (Maurice et al. 2011; Brum et al. 2016; Thingstad and Lignell 1997). In the nutrient-rich and microbial dense gut ecosystem, thus, one would expect a similar phage lytic strategy.

Yet the lysogenic life cycle seems to be prevalent in the gut, where upon infection the phage genome integrates in the bacterial host as a prophage, forming a phage-bacteria symbiont called lysogen. Markers of lysogeny have been observed in viral genomic and metagenomic data from healthy adults (Furuse et al. 1983; Letarov and Kulikov 2009; Reyes et al. 2010; Minot et al. 2011, 2013; Mirzaei and Maurice 2017; Beller and Matthijnssens 2019; Shkoporov and Hill 2019). Besides, the frequency of prophages in bacteria is positively correlated with the bacterial growth rate (Lauro et al. 2009; Touchon et al. 2016). This empirical observation in single cells aligns with having an increase of lysogeny in productive environments like the gut (Mirzaei and Maurice 2017; Kim and Bae 2018). Additionally, the virus-to-microbe ratio (VMR) in the gut is significantly lower than in environments with lower microbial densities (Knowles, Silveira, et al 2016; Wigington et al. 2016; Parikka et al. 2017). The combination of the factors mentioned above have led to the piggyback-the-winner (PtW) dynamics, which considers lysogeny as a substantial ecological strategy for phages in high microbial density environments (Knowles et al. 2016; Silveira and Rohwer 2016).

The prevalence of lysogeny in gut environments contrast with observations from marine studies over the years, where lysogeny had been characterized as an ecological strategy of phages to survive situations at limiting growth conditions (Weinbauer 2004). Prophages integrate in poor growth conditions and remain dormant until resources improve. Then the lytic pathway activates the production of new phage particles (Jiang and Paul 1998; Wilcox and Fuhrman 1994; Payet and Suttle 2013; Maurice et al. 2009; Paul and Weinbauer 2010; Maurice et al. 2011; Brum et al. 2016). This paradox of lysogeny at low versus high productive environments might stem from having overlooked the postive effects of lysogenic conversion in the bacterial host.

Prophages provide multiple positive attributes to the host (Brüssow et al. 2004; Howard-Varona et al. 2017). They confer immunity to the host cell against similar and dissimilar phages— superinfection exclusion mechanism—without necessarily compromising the host’s fitness in laboratory experiments (Bondy-Denomy et al. 2016; Mavrich and Hatfull 2019). They can also provide metabolic pathways that improve the host’s competitive edge in different conditions (Bossi et al. 2003; Edlin et al. 1975, 1977), supply regulatory proteins (Paul 2008), and promote the expression of promiscuous bacterial enzymes (Hultqvist et al. 2018). Superinfection exclusion provided by prophages also favors the acquisition of transducing virions that kill sensitive bacteria, increasing the gene transfer to lysogens (Haaber et al. 2016; Touchon et al. 2017). This includes antibiotic resistance genes (Colavecchio et al. 2017). Prophages also enhance the adaptation and fitness of lysogens by encoding bacterial virulence factors implicated in animal infection and immune response evasion (Costa et al. 2018; Ohnishi et al. 2001; Hayashi et al. 2001; Waldor and Mekalanos 1996; Fortier and Sekulovic 2013). In addition, the domestication of prophages in bacteria removes the capability of generating new viral particles while conserving phage genes (Bobay et al. 2014), a process linked to the evolutionary acquisition of important bacteria elements such as gene transfer agents (Lang et al. 2012), bacteriocins, and type VI secretion systems (Michel-Briand and Baysse 2002; Leiman et al. 2009).

Our hypothesis is that these beneficial aspects of lysogeny are particularly favorable at high production and bacterial concentration conditions. We propose that as microbial productivity increases, ecosystems transition through recurrent lysogenic-lytic cycles, as illustrated in Figure 1. In an established microbial community with lytic phage turnover, an increase in resources would favor the emergence of lysogens due to the superinfection exclusion defense mechanism and the other advantages conferred by prophages, in consonance with the piggyback-the-winner dynamics (Knowles et al. 2016; Silveira and Rohwer 2016). If the increase in resources is sustained, by virtue of the kill-the-winner dynamics (Thingstad et al. 2014; Våge et al. 2018), adapted virulent and temperate phages that can prey on abundant lysogens will rise, diversifying the community and increasing its richness (Martiny et al. 2014). As the microbial productivity and concentration increases, this recurrent lysogenic-lysis cycle would lead to a prophage ratchet, predicting an increase in the frequency of prophages in bacteria (Figure 1).

**Figure 1:**
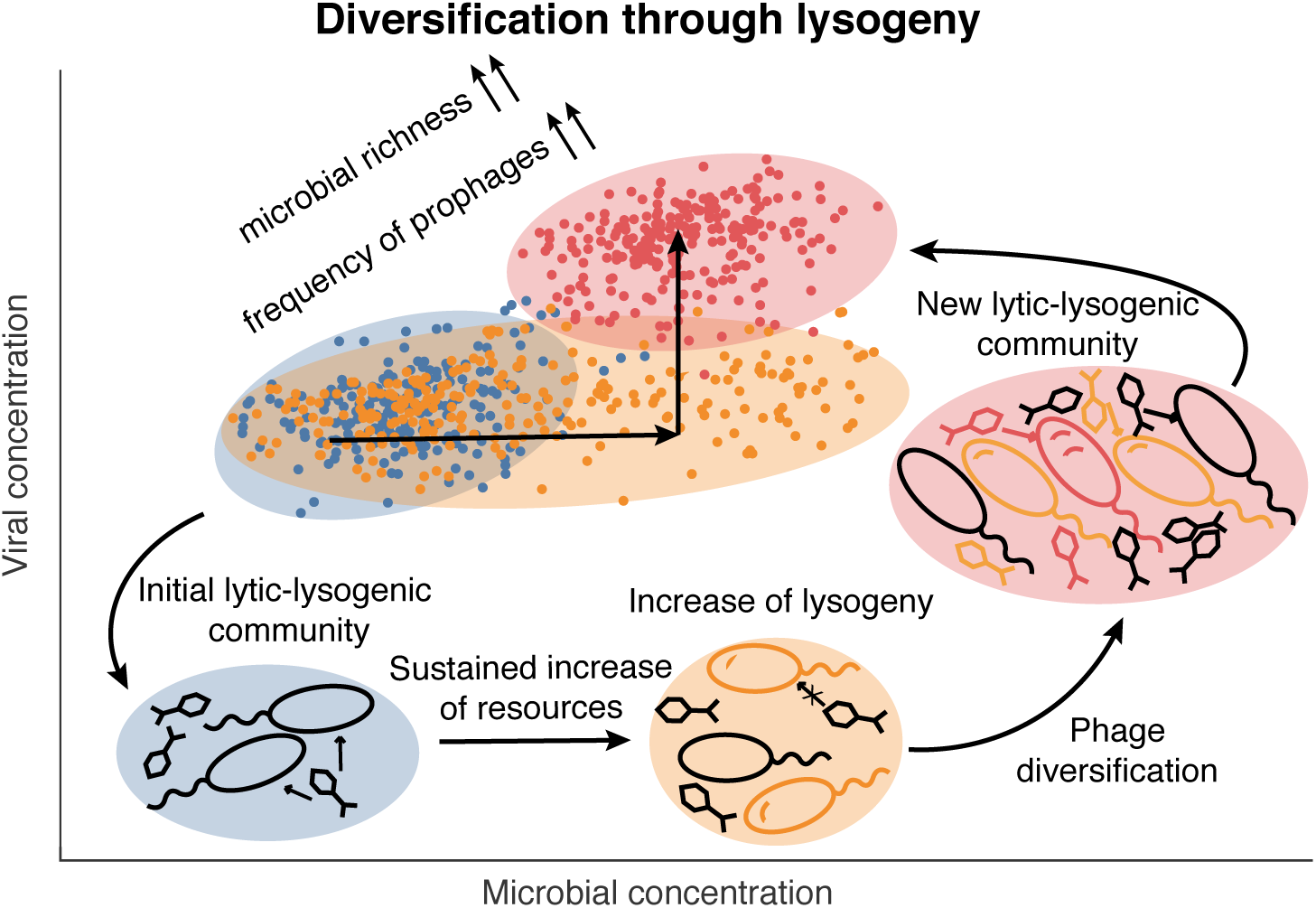
Diversification through lysogeny. The figure illustrates the central hypothesis of the research presented here. Upon increasing resources, an established viral-microbial community (blue) transitions to a new community (orange), where the microbial concentration increased due to the opportunistic growth of newly formed lysogens that escaped lytic phage top-down control by the superinfection exclusion mechanism. If the increase of resources is sustained, phages able to top-down control these newly form lysogens will eventually be selected and lead to a new mature viral-microbial community (red). This process increases the number of viral-host pairs (richness) and number of propahges per bacteria.

To test this hypothesis, we developed an ecological mathematical model with lytic and lysogenic phage-bacterial community compartments. This ecological model was coupled with a diversification model that estimated the accumulation of prophages in bacterial genomes. Using physiological values for the viral-bacterial traits extracted from the literature, the ecological model recovered viral and bacterial abundances observed across eleven ecosystems, and predicted a relationship between bacterial abundance and richness—defined from a viral predation standpoint. The diversification model predicted distributions of prophage per bacteria that agreed with abundances of prophages estimated bioinformatically in bacteria from aquatic environments and the human gut. It also provided predictions for the other ecosystems studied, which will further test the PtW-KtW cyclic framework introduced here.

## Methods

In this study, the terms phages and bacteria refer to the model. The terms viral-like-particles (VLPs) and microbial cells refer to environmental data. The terms viruses and microbes are used when discussing the model and environmental data.

### Lytic community compartment

The lytic compartment was modeled using a one-to-one network of phage and bacteria species. A bacterial species was defined ecologically based on being a prey of a different phage, and, for each community, the total number, *n*, of phage-bacteria species pairs was fixed. Phages in this community adopted the lytic cycle, without distinguishing between temperate and virulent phages. The model aimed to estimate the total phage and bacterial abundances in a community rather than the rank abundance of different taxonomic groups. The phage and bacterial life traits were approximated to be homogeneous in each community, but different communities had assigned different physiological values, which were explored stochastically (see the sampling section below). The net rate of each species in the community was given by

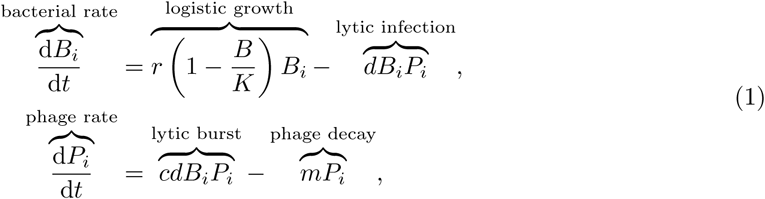

Here, *B*_*i*_ denoted each species of sensitive bacterium, and *P*_*i*_ each species of lytic phage. The index *i* identified each species and ranged from 1 to *n*, leading to 2*n* coupled equations per lytic community. The bacterial net production rate was the balance between the bacterial growth rate and the infection rate. In the bacterial growth term, the intrinsic growth rate *r* was reduced by a logistic factor that accounted for the total concentration of bacteria in the lytic community, 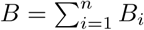, with respect to the carrying capacity, *K*. The functional response of phages praying on bacteria can introduce richer dynamics (Weitz and Dushoff 2008), but it is not usually considered in phage ecological models (Thingstad and Lignell 1997; Weitz et al. 2015) due to the lack of environmental parametrization values and its potential variability across different ecosystems (Hunsicker et al. 2011). The phage-bacteria infection mechanism was encoded as a standard mass action term, which has been reported to have a similar infection rate constant in different environments (Thingstad et al. 2014; Barr et al. 2015; Joiner et al. 2019). This reduced the number of parameters in the model. The infection rate was assumed to be proportional to the product of bacterial and viral species concentrations, that is, mass action for each phage-bacterial host pair, where *d* represented the infection rate constant. The viral net production was the balance of the lysis (burst size, *c*, times the infection rate) and viral decay (with a decay constant *m*).

The coexistence equilibrium concentrations for each agent in the community, 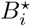 and 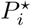, were obtained by solving analytically the steady-state of Eq. (1), that is, *dB*_*i*_*/dt* = 0 and *dP*_*i*_*/dt* = 0. Coexistence was required for the *n* species in the community, that is, 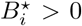 and 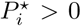. The stability conditions were obtained analytically using a linear approximation around the equilibrium values (Strogatz 2015). The Jacobian of the dynamical system was obtained at the coexistence equilibrium, and the determinant of the characteristic polynomial was transformed until extracting analytical expressions for all eigenvalues. The derivation is detailed in the Supplementary Information (S.1.1).

### Lysogenic community compartment

This community was formed by *n* temperate phage-lysogen pairs. Each lysogen had an active integrated prophage that could spontaneously induced, producing temperate phages. Thus this community was formed by *n* temperate phage-lysogen pairs. This compartment represented bacteria that had recently incorporated a prophage and was escaping the lytic top-down control in the community through the superinfection exclusion mechanism observed in laboratory experiments Brüssow et al. 2004. Except indicated otherwise, in the models the term lysogen refers to these escaping lysogenic community. The set of differential equations for the net production rate of escaping lysogens (*L*_*i*_) and temperate phages (*T*_*i*_) was given by

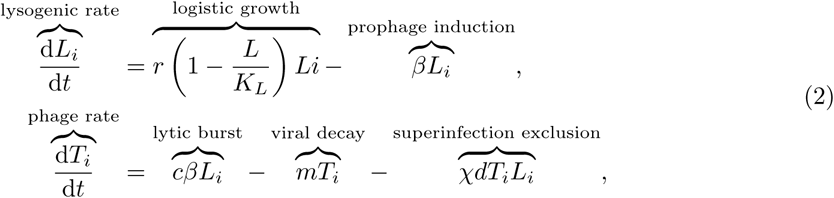

The index *i* identified each species and ranged from 1 to *n*, leading to 2*n* coupled equations as in the lytic community. The net production rate of lysogens was the balance between the lysogenic growth rate and the spontaneous prophage induction rate. The lysogenic growth was the intrinsic growth rate times a logistic term that accounted for the fraction of the total concentration of lysogens in the community, 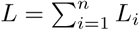, with respect to the carrying capacity in the lysogenic compartment, *K*_*L*_. The phage net production in the compartment was given by the production of temperate phages upon spontaneous prophage induction minus the phage decay and the removal of viable viruses due to superinfection exclusion. A value of χ = 1/2 accounted for a mix of surface defense mechanism with reversible phage-host binding (χ = 0 limit) and defense mechanism inactivating phage DNA (χ = 1 limit) Jasien 2017.

The equilibrium concentrations, 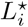 and 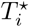, were obtained analytically by solving the algebraic equations that satisfied steady-state, *dL*_*i*_*/dt* =0 and *dT*_*i*_*/dt* = 0, for the coexistence regime, that is, 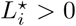 and 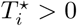 for the *n* species pairs. The stability conditions were obtained analytically using a linear approximation around the equilibrium values (Strogatz 2015). The Jacobian of the dynamical system was obtained at the coexistence equilibrium, and the determinant of the characteristic polynomial was transformed until extracting analytical expressions for all eigenvalues. The derivation is detailed in the Supplementary Information (S.1.2).

### Lytic-lysogenic coupling

The dynamics of the lytic-lysogenic model described in Eqs. (1) and (2) was studied for single species with additional terms accounting for direct interaction Jasien 2017. The superinfection exclusion mechanism led to a dominance of the lysogenic community. This was in agreement with laboratory experiments Chaudhry et al. 2019, but led to a relatively low phage concentration as well as phage-to-bacteria ratio compared to environmental data Wigingtonet al. 2016; Knowles et al. 2016; Parikka et al. 2017. To circumvent this issue, the lytic and lysogenic communities were treated in separated compartments, and the lytic community was given a preferential treatment. A given ecosystem was assumed to have an established lytic community with *n* phage-bacteria pairs. Sustaining this community in equilibrium required a carrying capacity *K > B*^⋆^. The remaining carrying capacity, Δ*K* = (*K − B*^⋆^), was assumed to provide potential resources for the community of lysogens to be established by escaping escaping top-down control due to the superinfection exclusion mechanism Brüssow et al. 2004. The available resources were assumed to promote the formation of new lysogens, and as they increased in abundance it was assumed that phages would evolve to apply top-down control, by virtue of the KtW motif (Vågeet al. 2018). The model presented here did not have the resolution to explore the arms race between this emerging lysogens and evolving phages. Instead, it assumed that the carrying capacity that escaping lysogens had access to grow, *K*_*L*_, was proportional to the remaining carrying capacity from the lytic compartment,

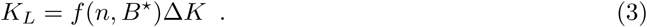

The *escaping lysogenic* factor, *f*, was established using physical arguments that led to

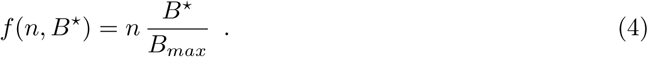

The arguments were as follows. First, it was assumed that new emerging lysogens would be more likely to form in a richer community. This was based on the lysogenic-lytic cycle hypothesis and the associated speciation through prophage integration. This lead to the proportionality term *n* in the escaping lysogenic factor above. Second, it was assumed that an increase in the concentration of top-down controlled bacteria in the lytic compartment would increase the encounter rate with temperate phages and promote the emergence of new lysogens. This led to the proportionality term *B*^⋆^. The escaping lysogenic factor must be unitless, so the contribution of the concentration needs to be normalized. This was done dividing the concentration *B*^⋆^ by the maximum packing concentration of bacteria *B*_*max*_ *≈* 10^12^ bacteria/mL. This was calculated by assuming a bacterial volume of 1 *µ*m3 (Milo and Phillips 2015). The total bacterial concentration was the sum of the bacteria in both compartments: The same applied to the total phage concentration:

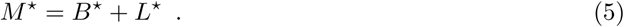

The same applied to the total phage concentration:

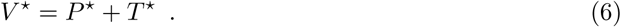

### Sampling of coexisting equilibria

Reference values for the intrinsic growth rate (*r*), infection rate (*d*), burst size (*c*), and decay rate (*m*) were obtained from Thingstad 2000 and Weitz et al. 2017, which published models that used established empirical values. The induction rate of prophages (*β*) was estimated from phage lambda (Rokney et al. 2008) and phage P1 (Rosner 1972). The range of carrying capacities were extrapolated from the microbial concentrations observed across ecosystems (Knowles et al. 2016). The reference value for each parameter can be found in Table 1. To account for environmental variability, each parameter was investigated for a range that spanned an order of magnitude higher and lower with respect to the reference value. The combination of parameters was explored using latin-hypercube sampling (LHS) (McKay et al. 1979; Weitz et al. 2017). For each given community, that is, a fix number of species *n*, 2500 equilibria points were sampled in three different scenarios: purely lytic, purely lysogenic, and lytic-lysogenic community. These equilibria included coexisting and non-coexisting solutions. The fraction of coexisting equilibria differed for different communities and scenarios, so 500 coexisting equilibria were randomly selected in each community for analysis. This protocol was applied to communities ranging from *n* =1 to 1000 phage-bacteria pairs.

**Table 1:** Parameters of the ecological model and reference empirical values. See the Methods section for the range explored for each parameter and literature sources.

| Parameter | Description | Environmental reference value |
| --- | --- | --- |
| $r$ | Bacterial growth constant | $\frac{1}{24} \text{ hr}^{-1}$ |
| $m$ | Viral decay constant | $\frac{1}{6} \text{ hr}^{-1}$ |
| $\beta$ | Induction rate of lysogen | $\frac{1}{24} \cdot 10^{-6} \text{ hr}^{-1}$ |
| $d$ | Infection rate | $10^{-8} \text{ mL bacteria}^{-1} \text{ hr}^{-1}$ |
| $c$ | Burst size | 20 |
| $\chi$ | Lysogen immunity | $\frac{1}{2}$ |
| $K$ | Carrying capacity | $10^4\text{--}10^{11} \text{ bacteria mL}^{-1}$ |

In the LHS, each parameter defined an orthogonal dimension on a hypercube. The range of each parameter was divided into an evenly spaced number of regions equal to the number of points sampled (2500 per fixed *n* and community model). Each sampling contained a value from a unique region of each parameter, and the value within this region was selected randomly from a uniform distribution. No partition was used more than once throughout the sampling process. All parameters were sampled in linear space except for the carrying capacity, which spanned over seven orders of magnitude (from 10^4^ to 10^11^ bacteria/ml) and was sampled in a logarithmic scale on base 10. For lytic communities with fixed richness, this sampling strategy was analogous to that introduced in Weitz et al. 2017.

A sensitivity analysis was applied to assess how the variation of the parameter values impacted the total concentration of bacteria (*M*) and phages (*V*). The analysis was performed around the mid-range reference value for the intrinsic growth rate (*r*), infection rate constant (*d*), decay rate constant (*m*), burst size (*c*), induction rate constant (*β*), superinfection exclusion factor (*x*), richness (*n*), and carrying capacity (*K*). Technical details are provided in the Supplementary Information, Section S.4 (Sensitivity analysis).

A variant of the model incorporating a grazing effect on bacteria was also explored. Grazers were introduced as a generic predator pressure on the bacterial population by the term (*-gB*_*i*_) in the population of bacteria. The grazer pressure rate constant, *g*, was assumed to be half of the bacterial replication rate *r/*2 (Fuhrman and Noble 1995; Thingstad and Lignell 1997). This term did not affect the results significantly. Grazers competed with phages, reducing their population abundance only in scenarios with relatively low carrying capacity. The main impact was the amount of percentage of sampled points that displayed stable coexistence, which dropped from 90% to 70%. To reduce the number of parameters in the model, the term was not included in the final version of the model.

### VLPs and microbial cell concentrations

Virus-like particles (VLP) and microbial cell counts were extracted from 22 studies covering eleven ecosystems as reported previously in Knowles et al. 2016. The meta-analysis included data from animal (Barr et al. 2013; Furlan 2009; Kim et al. 2011), coastal/estuarine (Bettarel et al. 2006; Bouvier and Maurice 2011; Hewson et al. 2001; Maurice et al. 2011, 2013; Parsons et al. 2015; Schapira et al. 2009), coral reef (Payet et al. 2014), deep ocean (Muck et al. 2014), drinking water (Rinta-Kanto et al. 2004), open ocean (Parsons et al. 2012), polar lakes (Laybourn-Parry et al. 2007; Lisle and Priscu 2004; Madan et al. 2005), sediment (Bettarel et al. 2006; Glud and Middelboer 2004; Mei and Danovaro 2004; Patten et al. 2008), soil (Amosse et al. 2013), soil pore water (Amosse et al. 2013), and temperate lake/river (Bouvier and Maurice 2011; Maurice et al. 2010) ecosystems. The studies used either flow cytometry or eplifuorescence to estimate cell and viral counts. Epifluorescence was used in 14 out of 22 studies for cell density (84% of the cell data) and 19 out of 22 studies for viral density (4% of the VLP data). The data on VLP and cell counts is provided in Source File 1. The raw image data were not available. The counts were obtained from tables and figures in the original manuscripts. The combined dataset was analyzed using the non-parametric method of smooth splines (*smooth.spline*) with cross-validation implemented by B. D. Ripley and Martin Maechler in the R language for statistical computing (Chambers et al. 1992; R Core Team 2018). Linear, least-squares regressions were calculated in R for logged VLPs and logged microbial concentrations as well as logged virus-to-microbe ratios (VMRs) and logged microbial concentrations. The logged data was in base 10. This analysis was applied to each ecosystem and all ecosystems combined. For the combined ecosystem analysis, the median values were used in the linear regression to reduce oversampling bias from individual ecosystems.

### Ecological model predictions

From the sampled communities in the model, linear statistical models were obtained for the richness, *n*, as a function of the median phage and bacterial concentrations using least-squares method. This statistical model was used to estimate the effective richness for each ecosystem studied using the median values for the VLPs and microbial cell abundances. For each ecosystem, the final richness was the average of the richness obtained from VLPs and microbial cell abundances. The percentage of bacteria escaping top-down control as lysogens was studied as a function of the community richness for the lytic-lysogenic model. A Hill function of order one (Michaelis-Menten) was fitted using a linear regression (least-squares method) on a double-reciprocal plot (Linewaver-Burk) to obtain a statistical model of the percentage of escaping lysogeny as a function of community richness. This statistical model was combined with the average richness obtained for each ecosystem to predict the percentage of the community escaping top-down control through lysogeny.

### Diversification model for the distribution of prophages per bacteria

A combinatoric model was developed to estimate the number of prophages per bacteria accumulated in different ecosystems. The integration of a prophage and its associated phenotypic advantages to the lysogen represent a substantial genetic variant with respect the original bacterium. Thus, in the diversification model, the integration of a prophage defined the formation of a new bacterial strain. At each diversification step, a newly evolved temperate phage was introduced in the community. Its integration as a prophage defined new bacterial strains. In the first evolutinary step (*z* = 1), the model contained a bacterial community composed of a single bacterial strain (*s* = 1) free of prophages (*q* = 0). In step two (*z* = 2), a temperate phage was introduced. Its integration as a prophage on an undetermined fraction of the initial bacaterial strain formed a community with two strains (*s* = 2): the original free-prophage bacterial strain (*q* = 0) and a new strain containing one prophage (*q* = 1). In the third diversification step (*z* = 3), a newly evolved temperate phage was added to the community. Its integration as a prophage promoted the formation of new strains. This led to three total strains (*s* = 3) and two alternative scenarios or states: in state 1, the prophage integrated in a fraction of the free prophage strain, leading to a community with a free-prophage strain (*q* = 0) and two strains with a different prophage each (two *q* =1 strains); in state 2, the prophage integrated in a fraction of the single-prophage strain from the prior diversification step, leading to a community with a free-prophage strain (*q* = 0), a one-prophage strain (*q* = 1), and a newly formed two-prophage strain (*q* = 2). This diversification process of generating strains was repeated subsequently for an arbitrary number of diversification steps, *z*, containing at each step *s* = *z* number of strains per state. A recursion relationship was derived to calculate the probability of having *q* prophages per bacteria in a system with *z* strains, *P* (*z, q*).

### Distribution of prophage per bacteria from low and high productive environments

The environmental distribution of prophages per bacteria was assessed from a low (marine surface) and high (human gut) productive environments. Phispy (version 3.2) (Akhter et al. 2012) was ran against a curated database of bacterial genomes from the Pathosystems Resource Integration Center (PATRIC). The process of curation focused on adding environmental annotation and removing plasmids, incomplete genomes, and metagenomes. A total of 83 genomes from marine surface bacteria and 750 from human gut were analyzed. Prophage genes were identified based on sequence similarity and genomic signatures as described previously (Akhter et al, 2012). A group of 30 or more prophage genes in a genome defined a single prophage in a bacterial genome. See Source Files 2 and 3.

### Diversification model predictions

The average number of prophages per bacteria obtained in marine surface and human gut environments was combined with the estimated richness from marine and animal gut communities to calibrate the prophage diversification and lytic-lysogenic community models. This facilitated the prediction of prophage frequency per bacteria in different ecosystems. The theoretical number of strains or diversification steps, *z*, was estimated by obtaining the theoretical average of prophages per bacteria, 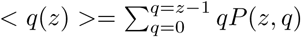, nearest to the average number of prophages obtained, respectively, for marine surface and human gut bacteria. The two theoretical diversification steps, *z*, were related with the effective number, *n*, of phage-bacteria pairs in marine surface and human gut assuming a power function model, *z*(*n*) = *an*^*b*^. This determined the values for *a* and *b*. The parameters obtained were *a* = 27.96 and *b* = 2.04. This power function model was applied to estimate the number of strains (diversification steps), *z*, as a function of the community richness, *n*, for each ecosystem studied. The number of strains (or steps) obtained determined the probability of number of prophages per bacteria, *P* (*z, q*), for the eleven ecosystems studied.

## Results

In the lytic community, Eq. (1), the equilibrium concentrations for each species of bacteria and phage were respectively, 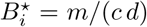 and 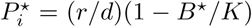, where *i* ranged from 1 to the mtotal number of species in the community *n*. The total bacterial and phage equilibrium concentrations were, respectively,

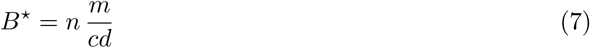

and

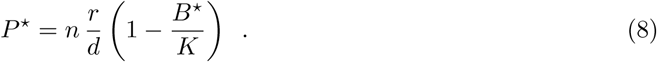

The total bacterial concentration, *B*^⋆^, was determined by the phage properties, that is, phage decay rate *m*, burst size *c*, and infection rate *d*. This was due to the top-down control applied by lytic phages. The bacterial population was also proportional to the richness of phage-bacteria pairs in the community *n*. The total phage concentration, *P*^⋆^, was a function of the bacterial growth rate *r*, the infection rate *d*, and the fraction of underutilized resources in the lytic community (1 − *B*^⋆^*/K*). The total phage concentration had a quadratic dependence on the phage-bacterial host pair richness, *n*. The community richness that maximized the concentration of phages was 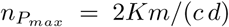. The analytical expressions for the 2*n −* 2 eigenvalues derived from the linear stability analysis indicated that the total phage and bacteria concentrations were stable for carrying capacities larger than the total bacterial population, *K > B*^⋆^ (see Supplementary section S.1.1). The individual species displayed an oscillatory behavior with an angular frequency 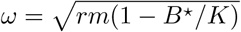. This was a consequence of having multiple one-to-one phage-bacteria pairs sharing the same resources, that is, carrying capacity, *K*.

In the lysogenic community, Eq. (2), the equilibrium distribution of lysogens was determined by the initial concentrations of lysogenic species, *L*_*i*_(*t*)*/L*(*t*)= *L*_*i*_(0)*/L*(0). The total concentration of lysogens was

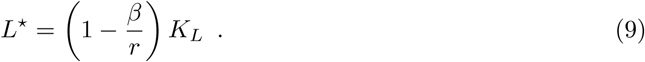

The total concentration of lysogens was proportional to the carrying capacity in the lysogenic compartment, *K*_*L*_, and it was reduced by the fraction of spontaneous prophage induction per replication (*β/r*). Thus, contrary to bacteria in the lytic compartment, the lysogenic community displayed a bottom-up control. This was a consequence of the superinfection exclusion mechanism, which removed the lytic predator pressure on the lysogens. The concentration of temperate phage species in the lysogenic community was 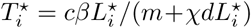. This led to a total concentration of temperate phages, *T*^⋆^, as a function of the distribution of lysogenic species. But the concentration of temperate phages generated from the lysogenic community had a small contribution to the pool of phages due to the small spontaneous induction rate. Two extreme cases illustrate this finding. A homogeneous distribution of lysogens, 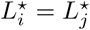, led to the total temperate phage concentration *T*^⋆^ = *cβL*^⋆^/(*mn* + *χdL*^⋆^). A community dominated by a single lysogenic species, 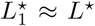 and 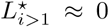, led to *T*^⋆^ = *cβL*^⋆^/(*m* + *χdL*^⋆^). Both expressions were Hill functions of order one with respect the lysogenic community, reaching the same plateau as the population of lysogens increased, regardless of the rank abundance of lysogenic species in the community,

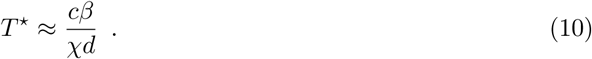

For the reference values given in Table 1, this was *T*^⋆^ *∼* 2 · 10^2^ phage/ml. Thus, the contribution of temperate phages from the active lysogenic community was small, and the choice of the specific distribution of the lysogenic community did not affect the results of the model significantly. The stability analysis was performed for the most general case, that is, with no specific distribution of lysogens (see Supplementary section S.1.2). There was *n* negative eigenvalues and *n −* 1 null eigenvalues, and the total concentration of lysogens remain stable for spontatenous induction rates smaller than the bacterial growth rate, *β < r*, which was the case for the empirical values, Table 1.

In the lytic-lysogenic community model, the total bacterial (*M*^⋆^) and phage concentrations (*V* ^⋆^), which combined the lytic and lysogenic communities sampled stochastically, covered, respectively, the range 10^4^–10^10^ bacteria/ml and 10^5^–10^10^ phages/ml, in consonance with the ranged observed across the eleven ecosystems studied (Figures 2**a** and 2**b**). The phage concentration increased sublinearly as a function of the bacterial concentration with an exponent *α* = 0.79 *±* 0.04 (SE) (R^2^ = 0.99, p *<* 0.001, *n*_*p*_ = 7), in agreement with the exponent observed environmentally (*α* = 0.84 *±* 0.11 (SE), R^2^ = 0.86, p *<* 0.001, *n*_*p*_ = 11). This power-function relating viral and microbial abundances was a first order approximation. A more refined statistical analyses presented in the Supplementary Information indicated that the exponent of the viral-microbial relationship becomes smaller with microbial concentration for both the model and the environmental data (Figures S.1, S.2, and S.3).

**Figure 2:**
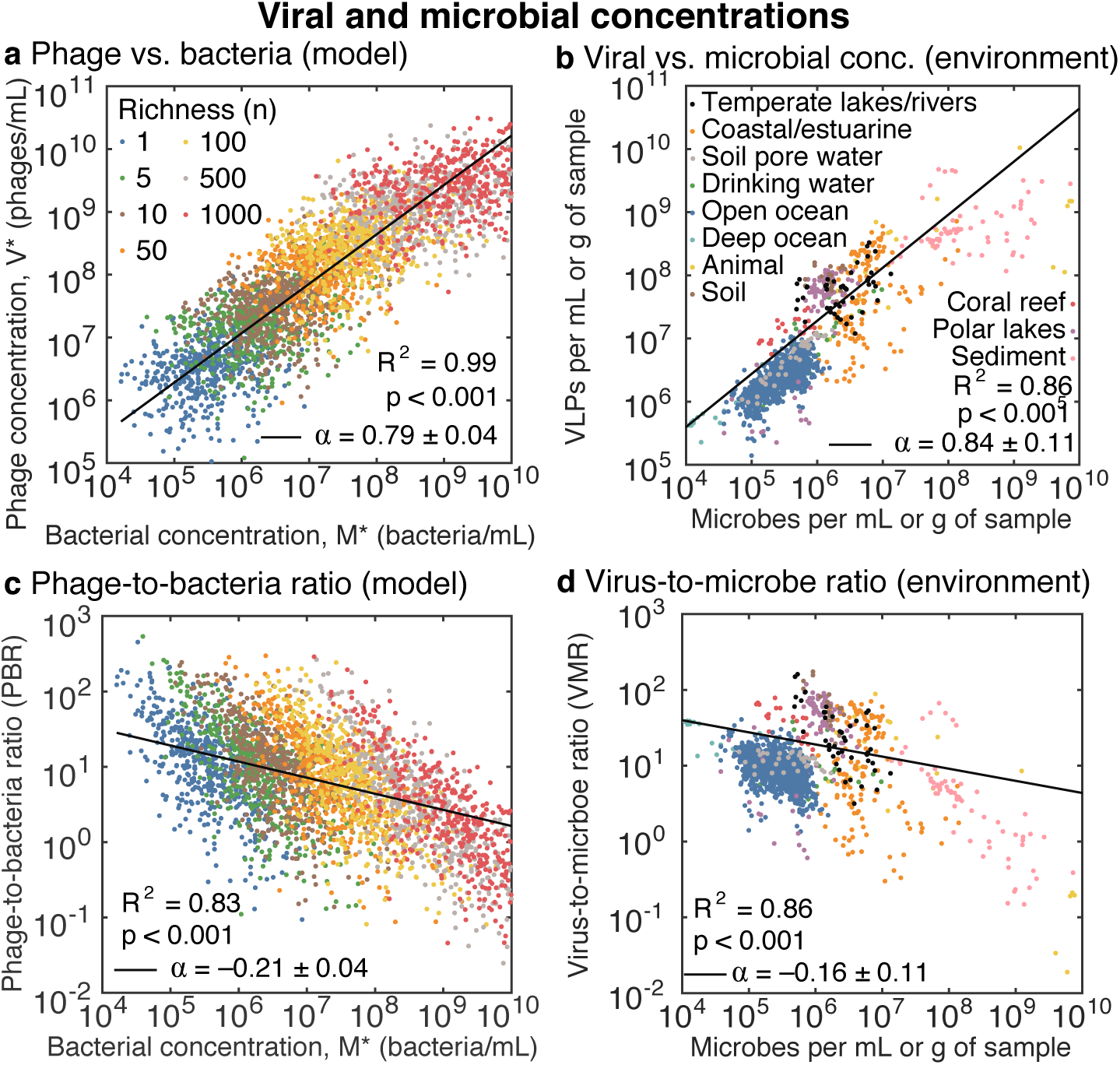
Viral and microbial concentrations. **a**, Phage equilibrium concentrations versus bacterial concentrations in the ecological lytic-lysogenic model. The colors distinguish communities with different richness, that is, number of bacteria species or phage-bacteria host pairs. **b**, Environmental concentration of virus-like particles (VLPs) versus microbial concentrations from eleven ecosystems (see legend for colors). Each ecosystems is plotted separately in Figure S.4. **c**, Phage-to-bacteria (PBR) versus bacterial concentration obtained in the model (same color coding as in panel **a**). **d**, Environmental virus-to-microbe ratio (VMR) versus microbial concentration for the ecosystems listed in panel **b. a-d.** The solid lines correspond to the linear regression of the logged data. The slope (*α ±* SE), p-value (*α* ≠ 0), and the coefficient of determination (*R*^2^) are displayed.

The phage-to-bacteria ratios (PBR) in the model ranged from 10^2^ to 10 ^*−*2^ phage/bacterium (Figures 2**c**). This aligned with thevirus-to-microbe ratios (VMR) observed environmentally (Figure 2**d**). PBR decreased as a function of bacterial concentration following a power function with exponent *α*_*PBR*_ = −0.21 *±* 0.04 (SE) (R^2^ = 0.83, p *<* 0.001, *n*_*p*_ = 7), which was consistent with the trend of environmental virus-to-microbe ratios (*α*_*VMR*_ = −0.16 *±* 0.11 (SE), R^2^ = 0.86, p *<* 0.001). As expected, the trend in PBR (or VMR) was related with the phage-bacteria (or viral-microbial) exponent by *α*_*PBR*_ *≈ α* − 1, since PBR = *V /M*. In communities with fixed richness, the stochastic sampling produces locally a sublinear trend that is intensified as lysogeny increases (Figure S.1). It is worth noting that no parameters were fitted in the model, except to the values used for the minimum and maximum richness (*n*).

The sensitivity analysis indicated that richness (*n*) was the most relevant parameter in the model, with a sensitivity factor *S*_*M,n*_ *≈* 1.5 in the bacteria population and *S*_*V,n*_ *≈* 1 in the phage population (Figure S.6). This was followed by the infection rate constant (*d*), which displayed a sensitivity factor *S ≈* 1 in both populations. The decay rate (*m*) and burst size (*c*) (both phage traits) where also sensitive in the bacteria population (sensitive factor *S ≈* 1), while the growth rate affected the phage population (sensitive factor *S ≈* 1). The carrying capacity was the last relevant parameter in the bacteria population (sensitive factor *S ≈* 0.5). The other parameters did not have a relevant impact in the community (sensitivity factor, *S ≈* 0). For further details, see Supplementary Information S.4.

The purely lytic community model, lacking the lysogenic compartment, recovered similar ranges of phage and bacteria concentrations (Figure S.5**a**). As in the case of the lytic-lysogenic community model, this was due to the variation of richness, *n*, across communities. The phage concentration followed a linear relationship with the microbial concentration (*α* = 1.00 *±* 0.01 (SE), R^2^ = 1.00, p *<* 0.001). Accordingly, the average phage-to-bacteria (PBR) ratio was constant, PBR *≈* 10 (Figure S.5**b**), that is, it did not vary with respect the microbial concentration (*α*_*PBR*_ = 0.00 *±* 0.01 (SE), R^2^ = 0.02, p = 0.736), in disagreement with the trend observed environmentally for the virus-to-microbe ratio (Figure 2**b**). Lysogeny, thus, was necessary in the model to recover a sublinear relationship between phage and bacterial concentrations across ecosystems.

The percentage of lysogeny, *L*[%] in the lytic-lysogenic community increased as a function of the total bacterial concentration, *M*^⋆^, saturating near 10^10^ bacteria/ml (Figure 3**a**). The parameters obtained for the fitted Hill function of order one were *a* = 97.1 *±* 0.7 and *b* = (2.86 *±* 8) · 10^8^ bacteria/ml (see details in the caption of Figure 3). The model indicated that the median percentage of lysogeny increased abruptly at bacterial concentrations larger than 10^6^ bacteria/mL. In particular, the median percentage of lysogeny was 1% at *∼* 3.0 · 10^6^ bacteria/mL, 10% at *∼* 3.3 · 10^7^ bacteria/mL, 50% at *∼* 3.0 · 10^8^ bacteria/mL, and 90% at *∼* 3.6 · 10^9^ bacteria/mL.

**Figure 3:**
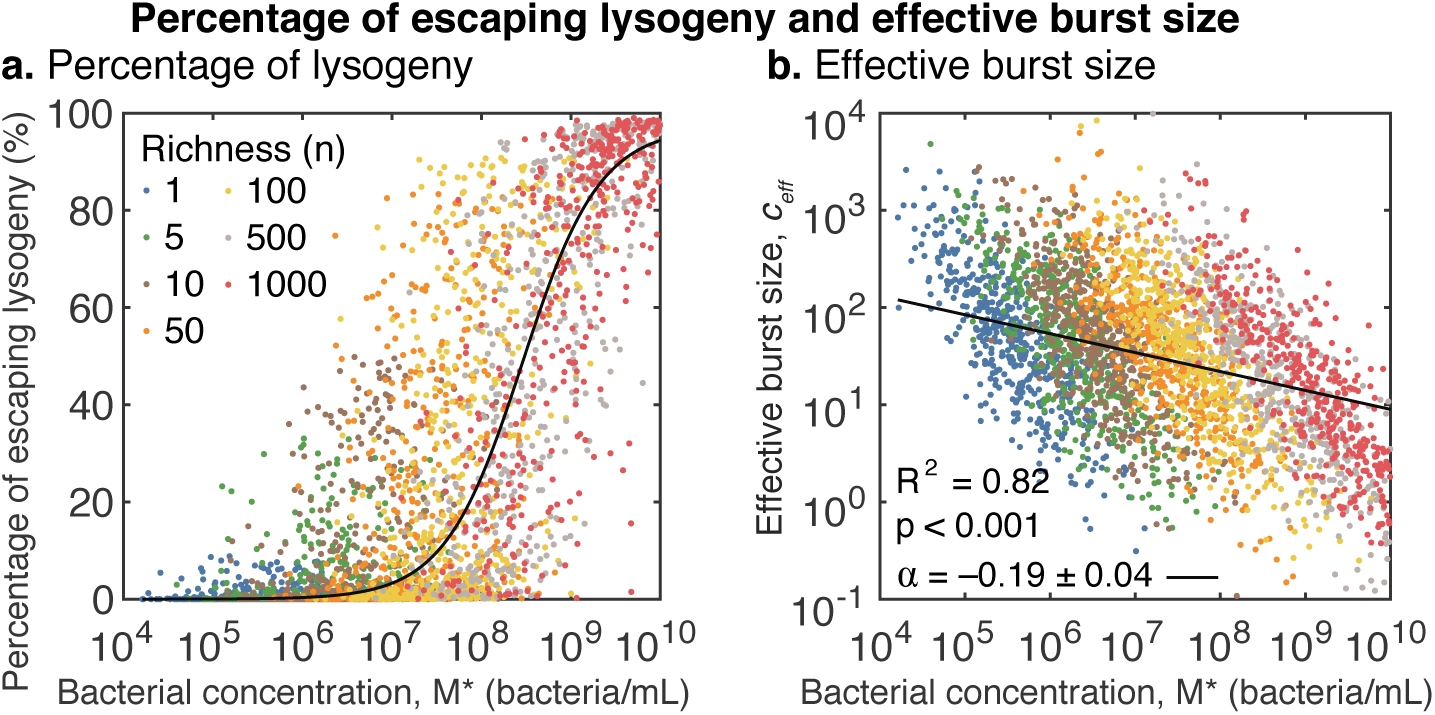
Percentage of lysogeny and effective burst size. **a**, Sampled percentage of active lysogeny as a function of bacterial concentration for the lytic-lysogenic community model. The solid line corresponds to the best fitted sigmoidal function *L*[%] = *aB*^⋆^/(*b* + *B*^⋆^) for the medians of the percentage of lysogeny, *L*[%], and total bacterial concentrations, *B*^⋆^, for each community, *n*. The parameters were obtained using the nonlinear least-squares Gauss-Newton algorithm implemented in the Nonlinear Least Squares (nls) function in R. The values obtained for the parameters corresponded to *a* = 97.1 *±* 0.7 and *b* = (2.86 *±* 8) · 10^8^ bacteria/ml. **b**, Effective burst size (*c*_*eff*_) as a function of bacterial concentration for the interpretation of the lytic-lysogenic community model as an effective single phage-bacteria community pair model. **a-b**, The colors correspond to communities with richness ranging from *n* =1 to *n* = 1000.

To facilitate the comparison with empirical data, the estimated phage-bacteria pair richness obtained for each ecosystem was expressed relative to the ecosystem with the lowest predicted richness, that is, deep ocean (Figure 4**a**). The relative richness obtained across ecosystems was *∼* 1 (deep ocean), *∼* 7 (open ocean), *∼* 12 (soil pore water), *∼* 16 (drinking water), *∼* 18 (coral reef), *∼* 64 (polar lakes), *∼* 85 (temperate lakes), *∼* 88 (coastal estuaries), *∼* 104 (soil), *∼* 248 (animal), and *∼* 978 (sediment). The phage-bacteria host pair richness of a community increased with the microbial and viral concentrations in ecosystems (Figure 2**b**).

**Figure 4:**
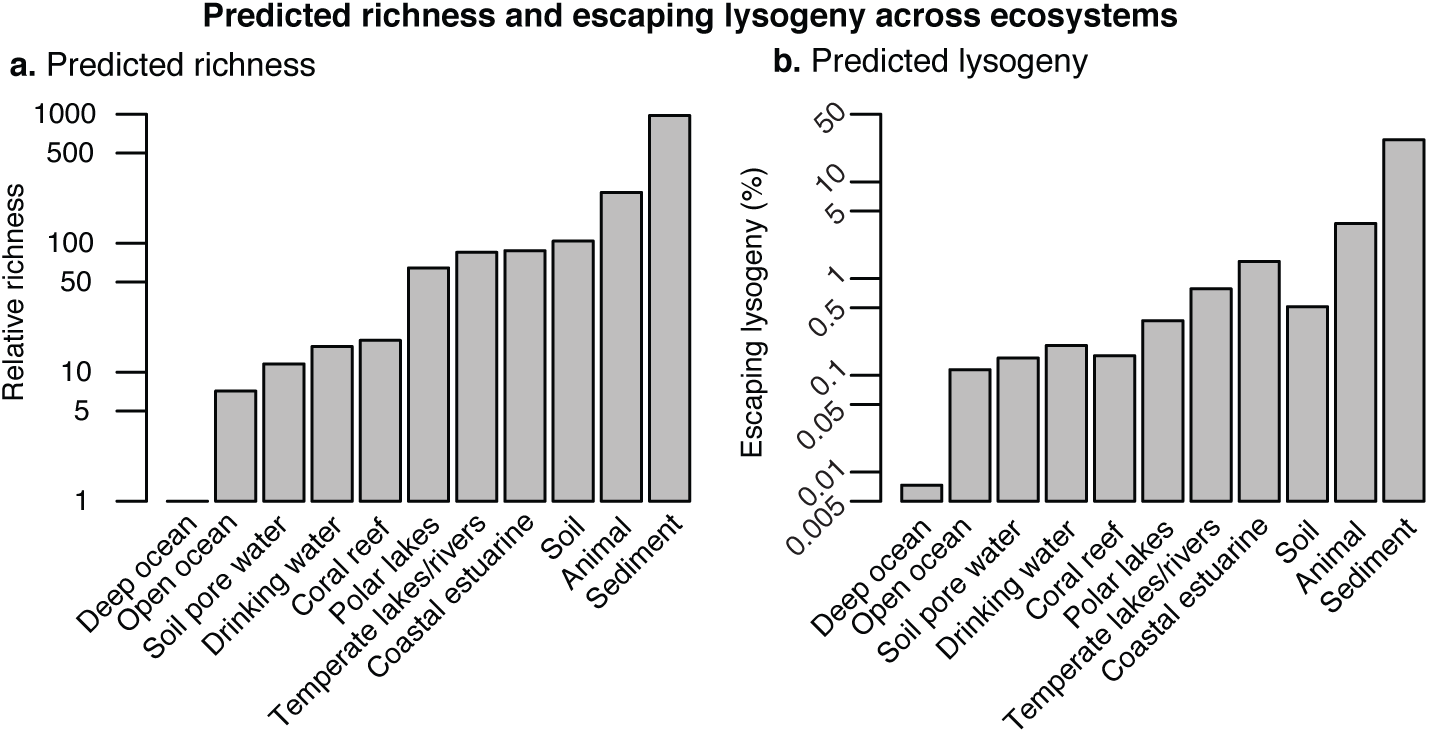
Predicted richness and escaping lysogeny across ecosystems. **a**, Relative community richness predicted for different ecosystems using the lytic-lysogenic community model. The values are given relative to the Deep Ocean data (lowest richness). **b**, Percentage of escaping lysogeny predicted by the model for the different ecosystems.

The percentage of escaping lysogens predicted across ecosystems was (Figure 4**b**) *∼*0.0073% (deep ocean), *∼*0.11% (open ocean), *∼*0.15% (soil pore water), *∼*0.20% (drinking water), *∼*0.16% (coral reef), *∼*0.37% (polar lakes), *∼*0.78% (temperate lakes), *∼*1.5% (coastal estuaries), *∼*0.5% (soil), *∼*3.7% (animal), and *∼*27% (sediment). The percentage of escaping lysogeny increased with richness, but some ecosystems, such as coral reefs and soils, displayed a relatively lower lysogeny with respect the contiguous ecosystems. This was due to their higher virus-to-microbe ratios.

In the diversification model, the number of bacterial strains with *q* prophages after *z* evolution steps (strains in the system) was related to the number of strains containing *q −* 1 prophages in the prior evolution step plus (*z −* 1)! times the number of bacterial strains having *q* in the prior step, that is, *R*(*z, q*)= *R*(*z −* 1,*q −* 1) + (*z −* 1)!*R*(*z −* 1, *q*). The total number of strains in the ensamble at step *z* was *z*! The probability of having *q* prophages per bacteria in a given evolutionary step was

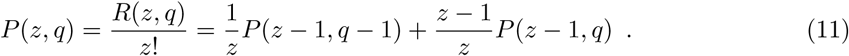

This determined the theoretical distribution of prophages per bacteria as a function of the total number of strains (evolutionary steps) *z*.

The empirical distributions of prophage per bacteria were obtained for superficial sea water (5 *±* 2 (SD) prophage/bacteria, 83 genomes) and human gut (14 *±* 5 (SD) prophage/bacteria, 750 genomes) (Figure 5**a**). The theoretical distributions were centered by fitting the prophage evolution generation number (*z*) to recover the average number of prophage per bacteria obtained empirically, leading to *z*_1_ = 182 (marine surface) and *z*_2_ = 1.41 · 10^6^ (human gut). The theoretical distributions of prophage per bacteria recovered the same distribution shape of the empirical data. By applying the effective richness (*n*) derived from the ecology model, the evolutionary model predicted the distributions of prophage per bacteria for each ecosystem (Figure 5**b**). The average number of prophages (*±* SD) per bacteria predicted for each ecosystem was (Figure 5**c**): open ocean (3.1 *±* 1.6), soil pore water (4.0 *±* 1.8), drinking water (5 *±* 2), coral reef (5 *±* 2), temperate lakes/rivers (8 *±* 3), polar lakes (8 *±* 3), coastal/estuarine (8 *±* 3), soil (9 *±* 3), animal (10 *±* 3), and sediment (13 *±* 4).

**Figure 5:**
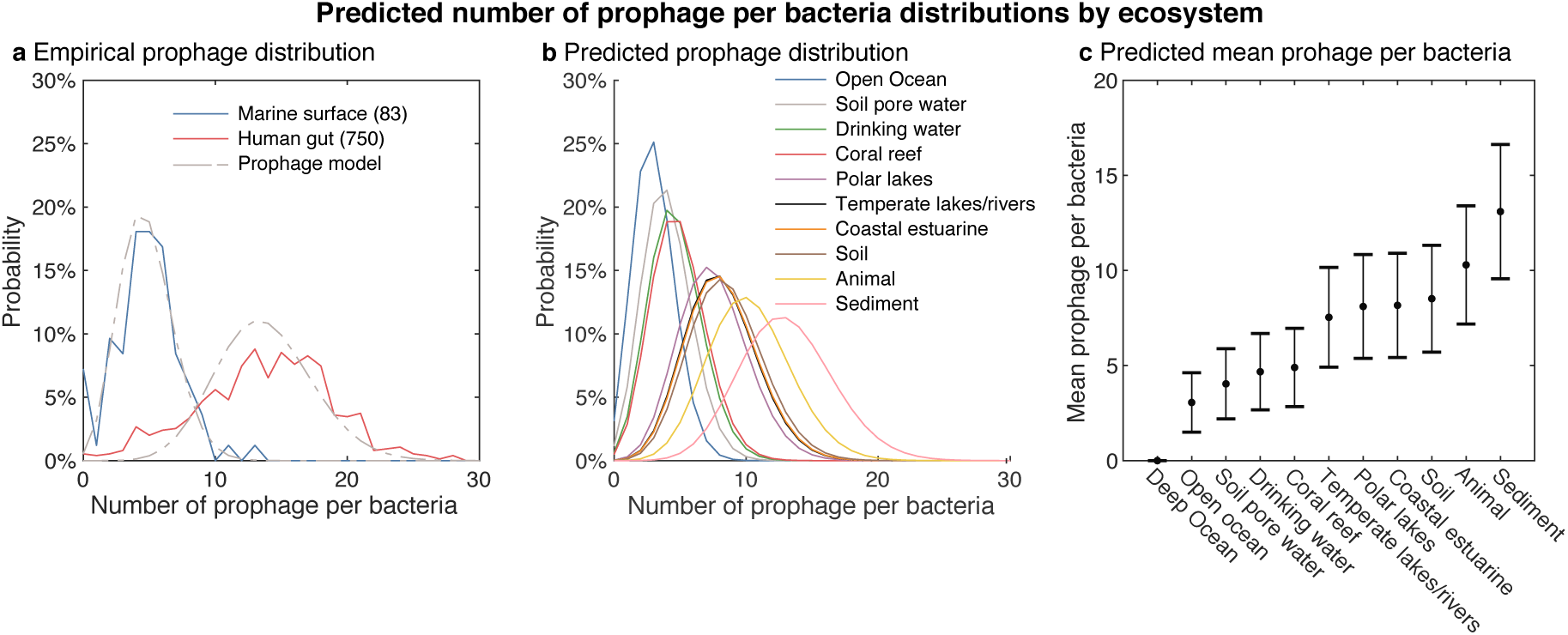
Evolutionary model predictions. **a**, Percentage of number of prophage per bacteria found in marine surface ecosystems (blue line) and human gut samples (red line), as well as the prophage distributions obtained from the prophage accumulation evolutionary model (grey dashed line). **b**, Predicted probability distributions for the number of prophages per bacteria in each ecosystem using the prophage accumulation evolutionary model. **c**, Mean number of prophage per bacteria predicted in each ecosystem. The error bars correspond to the associated standard deviations.

Finally, the lytic-lysogenic community model was then reinterpreted using an effective single lytic virus-host community with variable burst size. This aimed to faciliate the comparison of the lytic-lysogenic community model with the original PtW model (Knowles et al. 2016), which treated lysogeny implicitely in a single phage-bacteria species system by reducing burst size as a function of eutrophic conditions. The equilibrium values and stability of the lytic compartment were recovered by renormalizing the infection rate by the underlying community richness: *d → d/n* (see Eq. S.17). Note that the absence of the explicit phage-bacteria network removed the oscillatory modes obtained in the explicit lytic community. The lysogenic compartment was absorbed by the effective burst size *c*_*eff*_ = (*d/n*)*/m*(*B*^⋆^ + *L*^⋆^). This led to an effective burst size that decreased with bacterial concentration with a power law function with exponent *α* = −0.19 *±* 0.04 (SE) (R^2^ = 0.82, p *<* 0.001) (Figure 3**b**).

## Discussion

Over decades, lysogeny has been interpreted as an ecological strategy for phages to find refuge in their bacterial hosts during starvation and low production conditions (Weinbauer 2004; Paul and Weinbauer 2010; Maurice et al. 2009; Brum et al. 2016). The human gut and other host-associated microbiomes, however, are dominated by lysogenic bacteria and temperate phages despite being highly productive and microbial-dense environments (Minot et al. 2011; Reyes et al. 2010; Minot et al. 2013; Mirzaei and Maurice 2017; Kim and Bae 2018). To explain this paradox, here we hypothesized that the benefits acquired by the bacterial host during phage integration favor the emergence of lysogens at high microbial abundances. It is propose that lysogens escape the lytic top-down control of phages, and eventually phages evolve to lytically infect the escaping lysogens, forming a lysogeny-lysis cycle. The recurrence of this lysogeny and lysis cycle as resources increase lead to a diversification of the viral and microbial communities (Figure 1). This PtW-KtW cyclic hypothesis was tested by coupling a lytic-lysogenic ecological model and a prophage acquisition evolutionary model.

In the ecological model, lysogeny increased as a function of resources, density, and richness (Figure 3). The model recovered microbial and viral concentrations observed empirically across eleven ecosystems, ranging from 10^4^ cells/ml and 10^5^ phages/ml to 10^10^ cells/ml and 10^10^ phages/ml (Wigington et al. 2016; Knowles et al. 2016; Parikka et al. 2017). It also recovered the empirical sublinear trend between viral and microbial concentrations (exponent *∼* 0.7). In the model, the fraction of lysogens increased from 1% at *∼* 10^6^ bacteria/mL to 90% at *∼* 10^9^ bacteria/mL. This was a consequence of lysogenic bacteria escaping top-down control through superinfection exclusion (Brüssow et al. 2004; Touchon et al. 2017). Thus, in environments with *∼* 10^6^ bacteria/mL, the signal associated to lysogeny is predicted to be weak, in consonance with metagenomic observations in marine systems (Knowles et al. 2016; Luo et al. 2020). In environments with higher microbial density, instead, the lysogeny signal is predicted to be strong, a result consistent with observations in animal gut (Minot et al. 2011; Reyes et al. 2010; Minot et al. 2013; Mirzaei and Maurice 2017; Kim and Bae 2018).

Richness was defined as the number of phage-bacterial host pairs in the community richness, *n*. The increase in richness was associated to an increase of phage and bacteria concentrations. This is consistent with genomic data across gradients of microbial abundance, showing an increase of phage-host pair diversification (Coutinho et al. 2019). The purely lytic community model recovered similar ranges of phage and bacterial concentrations (Figure S.5a). In this scenario, however, the viral concentration across communities with different richness displayed a linear trend (exponent *∼* 1) with a constant average virus-to-microbe ratio of *∼* 10 (Figure S.5b)—in disagreement with the sublinear trend observed environmentally (Figure 2). For communities with fixed richness, however, the lytic model displayed a sublinear trend. A prior lytic model with fixed richness showed that this is a consequence of the stochastic sampling of the phage-bacteria traits (Weitz et al. 2017). The purely lytic community model, thus, displays a Simpson’s paradox, where the local sublinear trend observed in communities with fixed richness differs from the global trend when comparing communities across richness (Simpson 1951; Knowles et al. 2017). This paradox is a consequence of both viral and microbial concentrations scaling linearly with richness, Eqs. (8) and (7).

At each step of the diversification model, an evolved temperate phage was assumed to be able to integrate as a prophage, defining new bacterial strains in the community. The average number of prophages per bacteria increased with the number of diversification cycles, in consonance with the distributions of prophages observed in the genomes of marine and human gut bacteria (Figure 5a). The calibration of the model using the mean prophage number per bacteria in these two ecosystems predicted 10,000 times more PtW-KtW cycles in the human gut than in marine surface communities, consistent with the increase in the frequency of prophages as a function of the environment productivity (Lauro et al. 2009; Lee and Patterson 2002; Bakenhus et al. 2017; Touchon et al. 2016). The microbial community colonization of the human gut in early childhood is extremely dynamic, showing sequential changes in dominant phage and bacteria (Breitbart et al. 2008; Sharon et al. 2013; Lim et al. 2015; Mirzaei and Maurice 2017). The PtW-KtW cycles introduced here would eventually lead to the stable, richer, and lysogenic communities observed in adulthood (Minot et al. 2013; Reyes et al. 2010; Shkoporov and Hill 2019). The predicted distribution of prophages per bacteria in the other ecosystems studied can be tested emperically in future work (Figures 5b and 5c).

The accuracy of the model could be improved by adding the molecular mechanisms leading to the formation of new lysogens, which has been modeled at single strain-level but proven hard to incorporate into ecological scenarios (Steward and Levin 1984; Wang and Goldenfeld 2010; Maslov and Sneppen 2017; Wahl et al. 2018; Weitz et al. 2019; Chaudhry et al. 2019). The accuracy could be also improved by incorporating by incorporating specific predator and immune system pressure (Thingstad and Lignell 1997; Thingstad 2000; Winter et al. 2010; Dwayne et al. 2017; Caron et al. 2017; Talmy et al. 2019), variable energy sources (Thingstad and Lignell 1997; Weitz et al. 2015), and ecosystem-specific ranges for phage-bacteria traits and nested viral-microbial networks (Flores et al. 2011; Thingstad et al. 2014; Gao et al. 2017; Hendricks 1972; Kirchman 2016; De Paepe and Taddei 2006).

## Conclusion

The ecological-evolutionary community model introduced here assumed a recurrent cycle of lysogeny and lysis. Productive microbial conditions promoted lysogeny (piggyback-the-winner) and the selection of phages able to control lysogens protected by superinfection exclusion (kill-the-winner). The model recovered viral and microbial concentrations observed across 11 ecosystems and predicted an increase in the frequency of prophages in bacteria, consistent with observations in marine ecosystems and the mammalian gut. The prophage-acquisition ratchet explains the observation of high lysogeny in rich microbial environments.

## Supporting information

source file 1 dataset: viral_microbial abundances and methods

source file 2 prophages marine surface

source file 2 prophages marine surface

