## Supplemental Information for "Phage and bacteria diversification through a prophage acquisition ratchet"

Supplementary Information

### S.1 Ecological model

#### S.1.1 Lytic community model

The lytic community model developed was

$$\begin{aligned}\frac{\mathrm{d}B_i}{\mathrm{d}t} &= r\left(1-\frac{B}{K}\right)B_i - dB_iP_i \ , \\ \frac{\mathrm{d}P_i}{\mathrm{d}t} &= cdB_iP_i - mP_i \ ,\end{aligned} \tag{S.1}$$

where $i = 1, 2, \ldots, n$.

###### S.1.1.1 Equilibria

Setting $\frac{\mathrm{d}B_i}{\mathrm{d}t} = \frac{\mathrm{d}P_i}{\mathrm{d}t} = 0$ and solving for $B_i$ and $P_i$ reveals that there were three combinations of populations of bacteria and phage species for which the system was at equilibrium.

The first equilibrium was the trivial equilibrium, given by $(B_i^\star, P_i^\star) = (0, 0)$.

The second corresponded to not having viruses in the system, that is, $(B^\star, P^\star) = (K, 0)$. Here the community of bacteria reached the system's carrying capacity. In this case, the concentrations for individual species of bacteria were not explicit. The distribution of the bacteria species was determined from the ratio of $\frac{\mathrm{d}B_i}{\mathrm{d}t}$ and $\frac{\mathrm{d}B}{\mathrm{d}t}$, which led to the differential equation

$$\frac{\mathrm{d}B_i}{\mathrm{d}B} = \frac{r\left(1-\frac{B}{K}\right)B_i}{r\left(1-\frac{B}{K}\right)B} = \frac{B_i}{B} \ . \tag{S.2}$$

By separating variables and letting the initial condition of the system be $B_i(t_0) = B_{i0}$ and $B(t_0) = B_0 = \sum_{i=1}^{n} B_{i0}$, this led to the equation

$$\frac{B_i(t)}{B(t)} = \frac{B_{i0}}{B_0} \ . \tag{S.3}$$

This implied that the relative concentration of each bacteria species remained constant in time, that is, in the absence of phages, the initial conditions determined the distribution of bacteria species in the system.

The third case corresponded to the coexistence equilibrium:

$$B_i^\star = \frac{m}{cd} \ , \tag{S.4}$$

$$P_i^\star = \frac{r}{d}\left(1-\frac{mn}{cdK}\right) = \frac{r}{d}\left(1-n\frac{B_i^\star}{K}\right) . \tag{S.5}$$

The equilibrium of the total bacteria population was the sum over all species of bacteria:

$$B^{\star} = \sum_{i=1}^{n} \frac{m}{cd} = n\frac{m}{cd} = n\,B_i^{\star}\ . \tag{S.6}$$

Similarly, the expression for the population of the phage community was obtained by summing over all species of phage:

$$P^{\star} = \sum_{i=1}^{n} \frac{r}{d}\left(1 - \frac{mn}{cdK}\right) = \frac{rn}{d}\left(1 - \frac{mn}{cdK}\right) = \frac{rc}{m}B^{\star}\left(1 - \frac{B^{\star}}{K}\right). \tag{S.7}$$

Notice that the total concentration of phage was a concave down parabola with respect to the total concentration of bacteria. The maximum concentration was reached at $B^{\star} = K/2$, which defines implicitely the optimal number of species in the system that maximizes the phage population: $n_{opt} = cdK/(2m)$.

###### S.1.1.2 Stability

The Jacobian matrix of Eq. (S.1) was

$$J\Big|_{(B_i,P_i)} = \left(\begin{array}{ccccc|ccc}
r - \frac{rB}{K} - \frac{rB_1}{K} - dP_1 & -\frac{rB_1}{K} & \cdots & \cdots & -\frac{rB_1}{K} & -dB_1 & & 0 \\
-\frac{rB_2}{K} & \ddots & & & \vdots & & \ddots & \\
\vdots & & r - \frac{rB}{K} - \frac{rB_i}{K} - dP_i & & \vdots & & -dB_i & \\
\vdots & & & \ddots & -\frac{rB_{n-1}}{K} & & & \ddots \\
-\frac{rB_n}{K} & & & & r - \frac{rB}{K} - \frac{rB_n}{K} - dP_n & 0 & & -dB_n \\
\hline
& cdP_1 & & 0 & & cdB_1 - m & & 0 \\
& & \ddots & & & & \ddots & \\
& & cdP_i & & & & cdB_i - m & \\
& & & \ddots & & & & \ddots \\
& 0 & & cdP_n & & 0 & & cdB_n - m
\end{array}\right).$$

For the trivial equilibrium, $(B_i^{\star}, P_i^{\star}) = (0, 0)$, the Jacobian matrix reduced to

$$J\Big|_{(0,0)} = \left(\begin{array}{ccc|ccc}
\ddots & & 0 & & & \\
& r & & & \mathbf{0} & \\
0 & & \ddots & & & \\
\hline
& & & \ddots & & 0 \\
& \mathbf{0} & & & -m & \\
& & & 0 & & \ddots
\end{array}\right),$$

where the eigenvalues were $\lambda_i = r$ for $i = 1, \ldots, n$ and $\lambda_i = -m$ for $i = n+1, \ldots, 2n$. The Hartman-Grobman Theorem implied that the trivial equilibrium was always unstable (Wiggins

2003).

For the equilibrium with no phages, $(B^\star, P^\star) = (K, 0)$ the Jacobian matrix was

$$
J\Big|_{(B^\star,P^\star)} = \left(\begin{array}{ccccc|ccccc}
-\frac{-rB_1^\star}{K} & \frac{-rB_1^\star}{K} & \cdots & \cdots & \frac{-rB_1^\star}{K} & -dB_1^\star & & & & 0 \\
\frac{-rB_2^\star}{K} & \ddots & & & \vdots & & \ddots & & & \\
\vdots & & -\frac{rB_i^\star}{K} & & \vdots & & & -dB_i^\star & & \\
\vdots & & & \ddots & \frac{-rB_{n-1}^\star}{K} & & & & \ddots & \\
\frac{-rB_n^\star}{K} & & & & -\frac{rB_n^\star}{K} & 0 & & & & -dB_n^\star \\
\hline
 & & & & & cdB_1^\star - m & & & & 0 \\
 & & & & & & \ddots & & & \\
 & & 0 & & & & & cdB_i^\star - m & & \\
 & & & & & & & & \ddots & \\
 & & & & & 0 & & & & cdB_n^\star - m
\end{array}\right).
$$

The eigenvalues for this matrix satisfied the equation $\det\left(J\Big|_{(B_i^\star,P_i^\star)} - \lambda I\right) = 0$, that is,

$$
\left|\begin{array}{ccccc|ccccc}
\multicolumn{5}{c|}{A} & \multicolumn{5}{c}{B} \\
\hline
-\frac{-rB_1^\star}{K} - \lambda & \frac{-rB_1^\star}{K} & \cdots & \cdots & \frac{-rB_1^\star}{K} & -dB_1^\star & & & & 0 \\
\frac{-rB_2^\star}{K} & \ddots & & & \vdots & & \ddots & & & \\
\vdots & & \frac{-rB_i^\star}{K} - \lambda & & \vdots & & & -dB_i^\star & & \\
\vdots & & & \ddots & \frac{-rB_{n-1}^\star}{K} & & & & \ddots & \\
\frac{-rB_n^\star}{K} & & & & \frac{-rB_n^\star}{K} - \lambda & 0 & & & & -dB_n^\star \\
\hline
 & & & & & cdB_1^\star - m - \lambda & & & & 0 \\
 & & & & & & \ddots & & & \\
 & & 0 & & & & & cdB_i^\star - m - \lambda & & \\
 & & & & & & & & \ddots & \\
 & & & & & 0 & & & & cdB_n^\star - m - \lambda \\
\hline
\multicolumn{5}{c|}{C} & \multicolumn{5}{c}{D}
\end{array}\right| = 0
$$

Using properties of determinants of $2 \times 2$ block matrices, if $BC = CB$, then the above equation reduced to $\det(AD - BC) = 0$. Because $BC = CB = 0$ then $\det(A)\det(D) = 0$:

$$
\left|\begin{array}{ccccc}
\frac{-rB_1^\star}{K} - \lambda & \frac{-rB_1^\star}{K} & \cdots & \cdots & \frac{-rB_1^\star}{K} \\
\frac{-rB_2^\star}{K} & \ddots & & & \vdots \\
\vdots & & \frac{-rB_i^\star}{K} - \lambda & & \vdots \\
\vdots & & & \ddots & \frac{-rB_{n-1}^\star}{K} \\
\frac{-rB_n^\star}{K} & & & & \frac{-rB_n^\star}{K} - \lambda
\end{array}\right|
\left|\begin{array}{ccccc}
cdB_1^\star - m - \lambda & & & & 0 \\
 & \ddots & & & \\
 & & cdB_i^\star - m - \lambda & & \\
 & & & \ddots & \\
0 & & & & cdB_n^\star - m - \lambda
\end{array}\right| = 0
$$

The matrix on the right is a diagonal matrix and thus we can write the equation as

$$
\left|\begin{array}{ccccc}
\frac{-rB_1^\star}{K} - \lambda & \frac{-rB_1^\star}{K} & \cdots & \cdots & \frac{-rB_1^\star}{K} \\
\frac{-rB_2^\star}{K} & \ddots & & & \vdots \\
\vdots & & \frac{-rB_i^\star}{K} - \lambda & & \vdots \\
\vdots & & & \ddots & \frac{-rB_{n-1}^\star}{K} \\
\frac{-rB_n^\star}{K} & & & & -\frac{rB_n^\star}{K} - \lambda
\end{array}\right| \prod_{i=1}^{n} (cdB_i^\star - m - \lambda) = 0
$$

Adding multiples of rows and columns does not change the value of a determinant, therefore the operation $R_1 \rightarrow R_1 + \sum_{i=2}^{n} R_i$ was applied on the left determinant, where $R_i$ is the $i$th row. Keeping in mind that $\sum_{i=1}^{n} B_i^\star = B^\star = K$, this led to

$$\begin{vmatrix} -r-\lambda & -r-\lambda & \cdots & \cdots & -r-\lambda \\ \frac{-rB_2^\star}{K} & \ddots & & & \vdots \\ \vdots & & \frac{-rB_i^\star}{K}-\lambda & & \vdots \\ \vdots & & & \ddots & \frac{-rB_{n-1}^\star}{K} \\ \frac{-rB_n^\star}{K} & & & & -\frac{rB_n^\star}{K}-\lambda \end{vmatrix} \prod_{i=1}^{n}(cdB_i^\star - m - \lambda) \ = 0$$

$$(-r-\lambda)\begin{vmatrix} 1 & 1 & \cdots & \cdots & 1 \\ \frac{-rB_2}{K} & \ddots & & & \vdots \\ \vdots & & -\frac{rB_i}{K}-\lambda & & \vdots \\ \vdots & & & \ddots & -\frac{rB_{n-1}}{K} \\ \frac{-rB_n}{K} & & & & \frac{-rB_n}{K}-\lambda \end{vmatrix} \prod_{i=1}^{n}(cdB_i - m - \lambda) \ = 0$$

Then, the operation $C_i = C_i - C_{i-1}$ for the columns where $i = 2, \ldots, n$ led to

$$(-r-\lambda)\begin{vmatrix} 1 & 0 & \cdots & \cdots & 0 \\ \frac{-rB_2}{K} & -\lambda & & & \vdots \\ \vdots & & \ddots & & \vdots \\ \vdots & & & \ddots & 0 \\ \frac{-rB_n}{K} & & & & -\lambda \end{vmatrix} \prod_{i=1}^{n}(cdB_i - m - \lambda) \ = 0$$

$$(-r-\lambda)\,\lambda^{n-1}\prod_{i=1}^{n}(cdB_i - m - \lambda) = 0$$

The $2n$ eigenvalues of this matrix were $\lambda = -r$, $n-1$ copies of $\lambda = 0$, and $\lambda = cdB_i - m$ for $i = 1, \ldots, n$. This equilibrium was non-hyperbolic, and the Hartman-Grobman Theorem could not be applied to determine its stability (Wiggins 2003).

For the coexistence equilibrium, $(B_i^\star, P_i^\star) = \left(\frac{m}{cd}, \frac{r}{d}\left(1 - \frac{mn}{cdK}\right)\right)$, using $B^\star = \frac{mn}{cdK}$ the Jacobian matrix was

$$J\big|_{(B_i^\star, P_i^\star)} = \left(\begin{array}{ccc|ccc} \ddots & & \frac{-rm}{cdK} & \ddots & & 0 \\ & \frac{-rm}{cdK} & & & -\frac{m}{c} & \\ \frac{-rm}{cdK} & & \ddots & 0 & & \ddots \\ \hline \ddots & & 0 & & & \\ & rc\left(1-\frac{mn}{cdK}\right) & & & 0 & \\ 0 & & \ddots & & & \end{array}\right).$$

The eigenvalues for this matrix satisfied the equation $\det\left(J\big|_{(B_i^\star, P_i^\star)} - \lambda I\right) = 0$, that is,

$$
\left|\begin{array}{ccc|ccc}
\multicolumn{3}{c|}{A} & \multicolumn{3}{c}{B} \\
\hline
\ddots & & \frac{-rm}{cdK} & \ddots & & 0 \\
 & \frac{-rm}{cdK} - \lambda & & & -\frac{m}{c} & \\
\frac{-rm}{cdK} & & \ddots & 0 & & \ddots \\
\hline
\ddots & & 0 & \ddots & & 0 \\
 & rc\left(1 - \frac{mn}{cdK}\right) & & & -\lambda & \\
0 & & \ddots & 0 & & \ddots \\
\hline
\multicolumn{3}{c|}{C} & \multicolumn{3}{c}{D}
\end{array}\right| = 0
$$

$B$ and $C$ were diagonal matrices, thus the properties of determinants of $2 \times 2$ block matrices were applied because $BC = CB$, leading to leading to $\det(AD - BC) = 0$:

$$
\begin{vmatrix}
\ddots & & \frac{rm}{cdK}\lambda \\
 & \lambda^2 + \frac{rm}{cdK}\lambda + rm\left(1 - \frac{mn}{cdK}\right) & \\
\frac{rm}{cdK}\lambda & & \ddots
\end{vmatrix} = 0 \ .
$$

The entries along the diagonal were all the same, and all the other entries were the same different value. Taking $\alpha = \lambda^2 + \frac{rm}{cdK}\lambda + rm\left(1 - \frac{mn}{cdK}\right)$ and $\beta = \frac{rm}{cdK}\lambda$, this led to

$$
\begin{vmatrix}
\alpha & \beta & & \cdots & \beta \\
\beta & \ddots & \ddots & & \vdots \\
\vdots & \ddots & \alpha & \ddots & \\
 & & \ddots & \ddots & \beta \\
\beta & & \cdots & \beta & \alpha
\end{vmatrix} = 0 \ .
$$

by applying the row operation $R_i = R_i - R_1$ for $i = 2, \ldots, n$, one obtained

$$
\begin{vmatrix}
\alpha & \beta & \cdots & & \beta \\
\beta - \alpha & \alpha - \beta & 0 & \cdots & 0 \\
 & 0 & \ddots & \ddots & \vdots \\
\vdots & \vdots & \ddots & \ddots & 0 \\
\beta - \alpha & 0 & \cdots & 0 & \alpha - \beta
\end{vmatrix} = 0 \ .
$$

Then by apply the column operation $C_1 = C_1 + \sum_{i=2}^{n} C_i$, this led to

$$\begin{vmatrix} \alpha + (n-1)\beta & \beta & \cdots & & \beta \\ 0 & \alpha - \beta & 0 & \cdots & 0 \\ & 0 & & \ddots & \vdots \\ \vdots & & \ddots & & 0 \\ 0 & \cdots & & 0 & \alpha - \beta \end{vmatrix} = 0 \ .$$

This upper triangular determinant was just the product of the diagonal:

$$(\alpha + (n-1)\beta)(\alpha - \beta)^{n-1} = 0 \ .$$

Returning to the actual values of $\alpha$ and $\beta$ gives:

$$\left(\lambda^2 + \frac{rmn}{cdK}\lambda + rm\left(1 - \frac{mn}{cdK}\right)\right)\left(\lambda^2 + rm\left(1 - \frac{mn}{cdK}\right)\right)^{n-1} = 0$$

The $2n$ eigenvalues of this matrix were

$$\lambda = -\frac{rB^\star}{2K} \pm \sqrt{\left(\frac{rB^\star}{2K}\right)^2 - rm\left(1 - \frac{B^\star}{K}\right)} \tag{S.8}$$

and $2n - 21$ copies of

$$\lambda = \pm i\sqrt{rm\left(1 - \frac{mn}{cdK}\right)} = \pm i\sqrt{rm\left(1 - \frac{B^\star}{K}\right)} \ .$$

Of these eigenvalues $2n - 2$ of them have $\mathrm{Re}(\lambda) = 0$, thus the coexistence equilibrium is non-hyperbolic and the Hartman-Grobman Theorem cannot be applied to determine its stability Wiggins (2003).

The standard process to analyze the stability of hyperbolic equilibria is to perform a center manifold reduction, but this was not strictly possible in all the equilibrium scenarios. Thus, a numerical approach was used to explore the conjecture that the linear stability analysis was valid. In particular, based on the eigenvalues obtained above, it could be conjectured that for $K < \frac{mn}{cd}$ the bacteria only equilibrium was stable and when $K > \frac{mn}{cd}$ the coexistence equilibrium was stable. To test this conjectured, MATLAB's ode45 solver was utilized. Simulations were run for $K = 10^5, 10^6, 10^7$, and $10^8$. For each value of $K$, $10^3$ parameter samples were generated using latin hypercube sampling. For each parameter sample Equation (S.1) was solved over a large time span ($t \in [0\ ,\ 5 \times 10^4]$ hours). The stability of the equilibrium was identical for all $n > 1$. A community with $n = 2$ was chosen as an initial case study. The eigenvalues corresponding to the coexistence equilibrium indicated that there was oscillatory dynamics in the system. To account for this the bacteria and phage community concentrations were averaged over a range of times in the interval.

A kernel probability distribution was then fit to these averages for each value of $K$. This confirmed that for the total populations of bacteria and phage, the bacteria only equilibrium was stable when $K < \frac{mn}{cd}$ and the coexistence equilibrium was stable when $K > \frac{mn}{cd}$.

##### S.1.2 Lysogenic Community Model

The lysogenic community model was given by

$$\begin{aligned} \frac{\mathrm{d}L_i}{\mathrm{d}t} &= r\left(1-\frac{L}{K}\right)L_i - \beta L_i \\ \frac{\mathrm{d}T_i}{\mathrm{d}t} &= c\beta L_i - mT_i - \chi d T_i L_i \end{aligned} \tag{S.9}$$

where $i = 1, 2, \ldots, n$.

###### S.1.2.1 Equilibria

Setting $\frac{\mathrm{d}L_i}{\mathrm{d}t} = \frac{\mathrm{d}T_i}{\mathrm{d}t} = 0$ revealed two possible equilibria. First the trivial equilibrium given by $(L_i^\star, T_i^\star) = (0, 0)$, and, second, the coexistence equilibrium given by

$$L^\star = \left(1-\frac{\beta}{r}\right)K \ . \tag{S.10}$$

In this case, there was no explicit equilibrium conditions found for the individual lysogen, but this could be inferred from the ratio of $\frac{\mathrm{d}L_i}{\mathrm{d}t}$ and $\frac{\mathrm{d}L}{\mathrm{d}t}$:

$$\frac{\mathrm{d}L_i}{\mathrm{d}L} = \frac{(r\left(1-\frac{L}{K}\right)-\beta)L_i}{(r\left(1-\frac{L}{K}\right)-\beta)L} = \frac{L_i}{L} \ . \tag{S.11}$$

Separating variables and letting the initial conditions be $L_i(t_0) = L_{i0}$ and $L(t_0) = L_0$, this led to

$$\frac{L_i(t)}{L(t)} = \frac{L_{i0}}{L_0} \ . \tag{S.12}$$

Thus, the relative population of lysogens was constant and depended on the initial distribution of species. The population of temperate phage species at the coexistence equilibrium was

$$T_i^\star = \frac{c\beta L_i^\star}{m + \chi d L_i^\star} \ . \tag{S.13}$$

The population of the temperate phage community cannot be determined explicitly because it depended on the distribution of the lysogen species. The particular case for a homogeneous population, $L_i^\star = L_j^\star$, led to $L_i^\star = L^{star}/n$, then the total concentration of temperate phages was

$$T^\star = \frac{c\beta L^\star}{m + \chi d L^\star/n} \ . \tag{S.14}$$

##### S.1.2.2 Stability

The Jacobian matrix of Equation (S.9) was

$$
J\Big|_{(L_i,T_i)} = \left(\begin{array}{ccccc|ccccc}
r(1-\frac{L}{K})-\frac{rL_1}{K}-\beta & -\frac{rL_1}{K} & \cdots\cdots\cdots & & -\frac{rL_1}{K} & & & & & \\
-\frac{rL_2}{K} & \ddots & & & -\frac{rL_2}{K} & & & & & \\
\vdots & & r(1-\frac{L}{K})-\frac{rL_i}{K}-\beta & & \vdots & & & \mathbf{0} & & \\
\vdots & & & \ddots & -\frac{rL_{n-1}}{K} & & & & & \\
-\frac{rL_n}{K} & & & & r(1-\frac{L}{K})-\frac{rL_n}{K}-\beta & & & & & \\
\hline
c\beta-\chi dT_1 & & & & 0 & -m-\chi dL_1 & & & & 0 \\
& \ddots & & & & & \ddots & & & \\
& & c\beta-\chi dT_i & & & & & -m-\chi dL_i & & \\
& & & \ddots & & & & & \ddots & \\
0 & & & & c\beta-\chi dT_n & 0 & & & & -m-\chi dL_n
\end{array}\right).
$$

For the trivial equilibrium, $(L_i^\star, T_i^\star) = (0,0)$, the Jacobian matrix reduced to

$$
J\Big|_{(0,0)} = \left(\begin{array}{ccc|ccc}
\ddots & & 0 & & & \\
& r-\beta & & & \mathbf{0} & \\
0 & & \ddots & & & \\
\hline
\ddots & & 0 & \ddots & & 0 \\
& c\beta & & & -m & \\
0 & & \ddots & 0 & & \ddots
\end{array}\right),
$$

and the eigenvalues were

$$
\begin{aligned}
\lambda_i &= r-\beta, \text{for } i = 1,\ldots,n \text{ and} \\
\lambda_i &= -m, \;\text{ for } i = n+1,\ldots,\; 2n
\end{aligned} \tag{S.15}
$$

For typical environmental values, Table 1, $r > \beta$, thus the first $n$ eigenvalues were positive. The application of the Hartman Grobman Theorem implied that the trivial equilibrium was always unstable (Wiggins 2003).

For the coexistence equilibrium, $(L_i^\star, T_i^\star) = \left(L_i^\star, \frac{c\beta L_i^\star}{m+\chi dL_i^\star}\right)$, using $L^\star = \left(1-\frac{\beta}{r}\right)K$ the Jacobian matrix reduced to

$$
J\big|_{(L_i^\star,T_i^\star)} = \left(\begin{array}{ccccc|ccccc}
\frac{-rL_1^\star}{K} & \cdots & & \cdots & \frac{-rL_1^\star}{K} & & & & & \\
\frac{-rL_2^\star}{K} & \ddots & & & \frac{-rL_2^\star}{K} & & & & & \\
 & & \frac{-rL_i^\star}{K} & & \vdots & & & 0 & & \\
\vdots & & & \ddots & & & & & & \\
\frac{-rL_n^\star}{K} & & & & \frac{-rL_n^\star}{K} & & & & & \\
\hline
c\beta - \frac{c\beta L_1^\star}{m+\chi dL_1^\star} & & & & 0 & -m-\chi dL_1 & & & & 0 \\
 & \ddots & & & & & \ddots & & & \\
 & & c\beta - \frac{c\beta L_i^\star}{m+\chi dL_i^\star} & & & & & -m-\chi dL_i & & \\
 & & & \ddots & & & & & \ddots & \\
0 & & & & c\beta - \frac{c\beta L_n^\star}{m+\chi dL_n^\star} & 0 & & & & -m-\chi dL_n
\end{array}\right),
$$

The eigenvalues were given by $\det\left(J\big|_{(L_i^\star,T_i^\star)} - \lambda I\right) = 0$. This matrix had $2n$ eigenvalues that satisfied

$$
\left|\begin{array}{ccccc|ccccc}
\frac{-rL_1^\star}{K} - \lambda & \frac{-rL_1^\star}{K} & \cdots & \cdots & \frac{-rL_1^\star}{K} & & & & & \\
\frac{-rL_2^\star}{K} & \ddots & & & \frac{-rL_2^\star}{K} & & & & & \\
 & & \frac{-rL_i^\star}{K} - \lambda & & \vdots & & & 0 & & \\
\vdots & & & \ddots & \frac{-rL_{n-1}^\star}{K} & & & & & \\
\frac{-rL_n^\star}{K} & & & & \frac{-rL_n^\star}{K} - \lambda & & & & & \\
\hline
c\beta - \frac{c\beta dL_1^\star}{m+dL_1^\star} & & & & 0 & -m-dL_1-\lambda & & & & 0 \\
 & \ddots & & & & & \ddots & & & \\
 & & c\beta - \frac{c\beta dL_i^\star}{m+dL_i^\star} & & & & & -m-dL_i-\lambda & & \\
 & & & \ddots & & & & & \ddots & \\
0 & & & & c\beta - \frac{c\beta dL_n^\star}{m+dL_n^\star} & 0 & & & & -m-dL_n-\lambda
\end{array}\right| = 0
$$

By using the aforementioned techniques to simplify the determinant of a $2 \times 2$ block matrix (because $BC = CB$), the determinant reduced to

$$\begin{vmatrix} \frac{-rL_1^\star}{K}-\lambda & \frac{-rL_1^\star}{K} & \cdots & \cdots & \frac{-rL_1^\star}{K} \\ \frac{-rL_2^\star}{K} & \ddots & & & \frac{-rL_2^\star}{K} \\ & & \frac{-rL_i^\star}{K}-\lambda & & \vdots \\ \vdots & & & \ddots & \frac{-rL_{n-1}^\star}{K} \\ \frac{-rL_n^\star}{K} & & & & \frac{-rL_n^\star}{K}-\lambda \end{vmatrix} \begin{vmatrix} -m-dL_1-\lambda & & & & 0 \\ & \ddots & & & \\ & & -m-dL_i-\lambda & & \\ & & & \ddots & \\ 0 & & & & -m-dL_n-\lambda \end{vmatrix} = 0$$

The second determinant in the above product was a diagonal matrix, so

$$\begin{vmatrix} \frac{-rL_1^\star}{K}-\lambda & \frac{-rL_1^\star}{K} & \cdots & \cdots & \frac{-rL_1^\star}{K} \\ \frac{-rL_2^\star}{K} & \ddots & & & \frac{-rL_2^\star}{K} \\ & & \frac{-rL_i^\star}{K}-\lambda & & \vdots \\ \vdots & & & \ddots & \frac{-rL_{n-1}^\star}{K} \\ \frac{-rL_n^\star}{K} & & & & \frac{-rL_n^\star}{K}-\lambda \end{vmatrix} (-m-dL_1^\star-\lambda)\cdots(-m-dL_i^\star-\lambda)\cdots(-m-dL_n^\star-\lambda) = 0$$

To extract the eigenvalues associated to the first determinant, the elementary row operation

$$R_1 = R_1 + \sum_2^n R_i$$

was applied combined with the constraint that $L^\star = \left(1 - \frac{\beta}{r}\right) K$,

$$\begin{vmatrix} \beta-r-\lambda & \cdots & \cdots & \beta-r-\lambda \\ -\frac{rL_2^\star}{K} & -\frac{rL_2^\star}{K}-\lambda & & -\frac{rL_2^\star}{K} \\ \vdots & \ddots & & \vdots \\ & & \ddots & \frac{-rL_{n-1}^\star}{K} \\ \frac{-rL_n^\star}{K} & & & -\frac{rL_n^\star}{K}-\lambda \end{vmatrix} \prod_{i=1}^{n}(-m-dL_i^\star-\lambda) = 0$$

By applying the elementary column operation $C_i = C_i - C_{i-1}$ for $i = 2, \ldots, n$, this resulted in

$$\begin{vmatrix} \beta-r-\lambda & 0 & \cdots & 0 \\ -\frac{rL_2^\star}{K} & -\lambda & & \vdots \\ \vdots & 0 & \ddots & 0 \\ \frac{-rL_n^\star}{K} & 0 & 0 & -\lambda \end{vmatrix} \prod_{i=1}^{n}(-m-dL_i^\star-\lambda) = 0$$

The first factor was a lower triangular determinant, thus

$$(\beta - r - \lambda)(-\lambda)^{n-1} \prod_{i=1}^{n}(-m-dL_i^\star-\lambda) = 0\,. \tag{S.16}$$

This led to the eigenvalues $\lambda = \beta - r$, $n-1$ copies of $\lambda = 0$, and $\lambda = -m - \chi dL_i^\star$ for $i = 1, \ldots, n$.

This equilibrium was non-hyperbolic and the Hartman Grobman Theorem cannot be applied to guarantee its stability. A numerical stability analysis similar to that completed in section S.1.1.2 was performed to double check the stability in the regions of interest.

#### S.2 Analysis of viral and microbial abundances

The outputs of the lytic-lysogenic model for communities ranging in richness from $n = 1$ to 1000 were merged and analyzed as a single dataset (Figure S.1**a**). The smooth spline had 6.8 degrees of freedom (Df), a smoothing parameter of 1.09 (spar), an associated lambda of $3.3 \cdot 10^{-2}$, and a penalized criterion (RSS) of 973.7. The spline displayed an initial trend with a smaller slope, in the region of $10^6$ bacteria/ml to $10^8$ bacteria/ml there is an increase in slope, with a change of concavity around $10^7$ bacteria/ml. At larger bacterial concentrations the slope decreases again. A similar analysis was applied to the environmental data. The viral and microbial abundances from the eleven ecosystems investigated were merged and analyzed as a single dataset (Figure S.1**b**). The smooth spline had 20 degrees of freedom (Df), a smoothing parameter of 0.97 (spar), a lambda of $6.9 \cdot 10^{-5}$, and a penalized criterion (RSS) of 159.7. From $10^4$ cells/ml to $10^6$ cells/ml the curve displayed a relatively linear trend. This was followed by a sudden shift to higher viral densities followed by oscillations that displayed a slower average increase of viral concentrations as microbial concentration increased. Thus, the smooth splines in both the model and the environmental data displayed a similar qualitative behavior, although the environmental data displayed additional degrees that gave rise to oscillations.

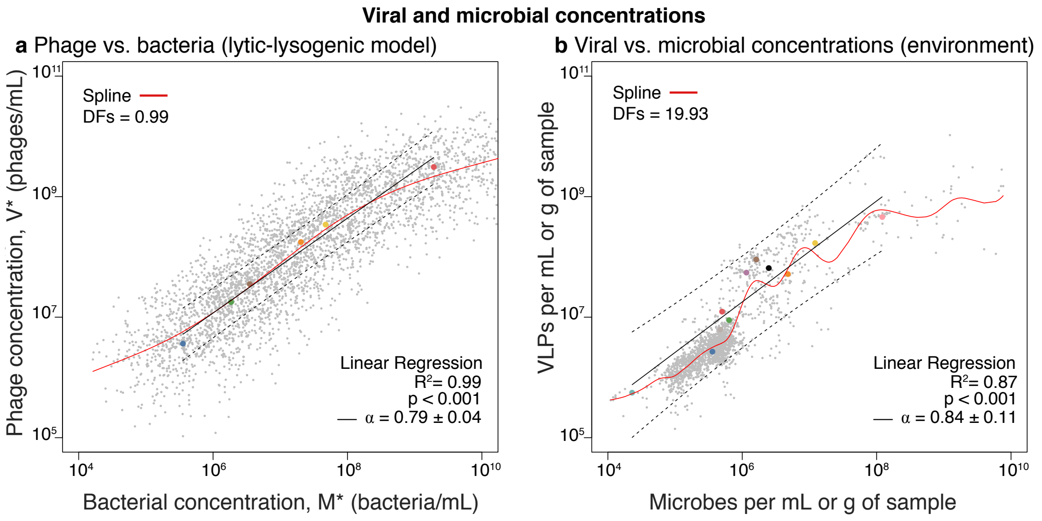


Figure S.1: **Viral and microbial abundance across ecosystems. a** Phage concentration as a function of bacteria concentration for communities with richness ranging from $n = 1$ to 1000. The grey dots are the outcomes for each sampling, while the colored circles correspond to the medians of each community with the same richness. The color code is equivalent to Figure 1**a**. **b** Viral-like particle (VLP) concentration as a function of microbial concentration across ecosystems. The grey dots are the outcomes of each sampling, while the colored circles correspond to the medians of each ecosystem. The color code is equivalent to Figure 1**b**. **a-b** The red line corresponds to a smooth spline, and the legend displays the associated degrees of freedom. The black line corresponds to a linear regression of the logarithmic data (base 10) for the medians. The legend displays the coefficient of determination ($R^2$), the p-value of the slope ($\alpha$), and the slope (with standard error). The dashed black lines represent the upper and lower predictions within the 95% level.

Next, the data of the lytic-lysogenic model was organized in groups of communities with the same richness ($n$). The logarithms in base 10 of the medians were analyzed using a linear model (that is, an implicit power function model). This led to a linear regression with intercept $\beta = 2.4 \pm 0.3$ (S.E), slope $\alpha = 0.79 \pm 0.04$ (p-value$= 8.9 \cdot 10^{-6}$), and coefficient of determination $R^2 = 0.99$ (Figure S.1**a**). The diagnostic of the residuals confirmed a slight non-linearity, but the values remained consistent with a normal distribution (Shapiro-Wilk test: W$= 0.90$, p-value $= 0.34$). The medians for each ecosystem in the environmental data were analyzed analogously (Figure S.1**b**). This led to a linear regression with intercept $\beta = 2.2 \pm 0.7$ (S.E), a slope $\alpha = 0.84 \pm 0.11$ (p-value$= 2.6 \cdot 10^{-5}$), and a coefficient of determination $R^2 = 0.87$. The diagnostic of the residuals indicated that the linear model was a good first approximation to quantify the trend in the data despite the oscillations observed in the smooth spline. The residuals did not display an apparent non linearity, and they were consistent with a normal distribution (Shapiro-Wilk test: W$= 0.98$, p-value $= 0.74$). Thus, the linear regression (log-log model) provides a reasonable first approximation to quantify the trends in the viral-microbial relationship, and the intercepts and slopes of the lytic-lysogenic model were quantiatively consistent with the environmental data.

A more refined analysis was explored by dividing the viral and microbial abundances in a set of low and high microbial concentrations (Figure S.2). The criteria to divide the change in trends observed in the smooth spline for the full datasets (Figure S.1). The data from the model was splitted at $10^7$ bacteria/ml (Figure S.2**a**). This was based on the change of concavity observed in the smooth spline (Figure S.1**a**). This also guaranteed an even distribution of communities within the low and high abundances. The linear regression of the subset of data with low microbial concentrations ($< 10^7$ bacteria/ml) led to a model with slope $\alpha = 0.71 \pm 0.02$ (p-value$< 2 \cdot 10^{-16}$) and a coefficient of determination $R^2 = 0.55$. The linear regression of the subset of data with high microbial concentrations ($> 10^7$ bacteria/ml) led to a model with slope $\alpha = 0.52 \pm 0.01$ (p-value$< 2 \cdot 10^{-16}$) and a coefficient of determination $R^2 = 0.52$. The environmental data was splitted at $10^6$ cells/ml (Figure S.2**b**). This was based on the sudden jump observed in the associated smooth spline at this concentration (Figure S.1**b**). As in the case of the model, this splitted led to an even distribution of ecosystems in each subset. The linear regression of the subset of data with low microbial concentrations ($< 10^6$ cells/ml) led to a model with slope $\alpha = 0.79 \pm 0.02$ (p-value$< 2 \cdot 10^{-16}$) and a coefficient of determination $R^2 = 0.49$. The linear regression of the subset of data with high microbial concentrations ($> 10^6$ cells/ml) led to a model with slope $\alpha = 0.52 \pm 0.03$ (p-value$< 2 \cdot 10^{-16}$) and a coefficient of determination $R^2 = 0.52$. Thus, for both the lytic-lysogenic community model and the environmental data, the slope (power function exponent) is steeper at low concentrations and reduces at higher microbial concentrations. The values of the slopes were similar for the lytic-lysogenic model and the environmental data. The discrepancy about the concentration splitting both regions was probably due to the generic use of phage-bacteria parameters in the lytic-lysogenic model, which are not specialized for specific ecosystems and could not capture variations between ecosystems in detail.

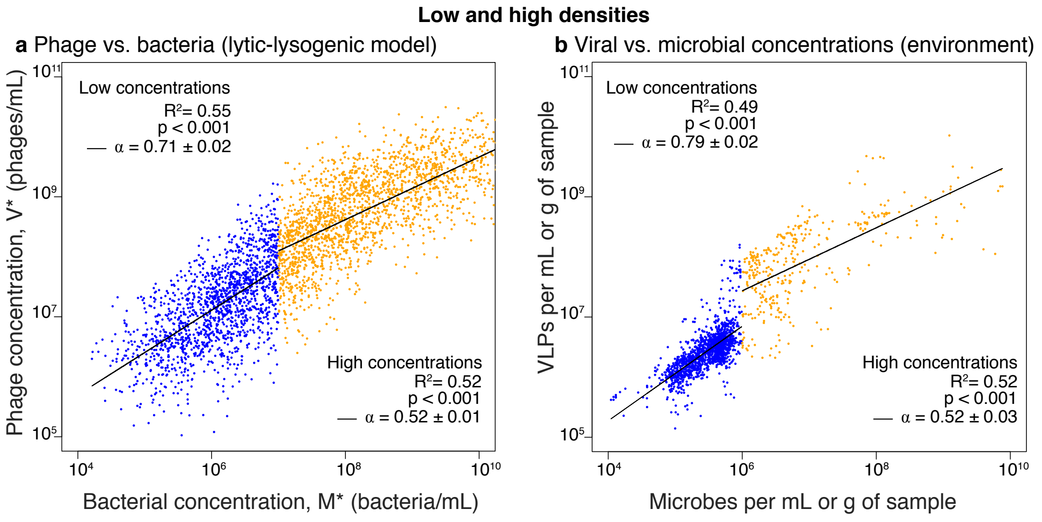


Figure S.2: **Low and high viral and microbial abundances.** **a** Phage concentration as a function of bacteria concentration for communities with richness ranging from $n = 1$ to 1000. The blue dots are the outcomes of samplings with bacterial concentrations lower than $10^7$ bacteria/ml. The orange dots correspond to cases with bacterial concentrations larger than $10^7$ bacteria/ml. **b** VLPs concentration as a function of microbial concentration for the eleven ecosystems studied. The blue dots are the outcomes of samplings with microbial concentrations lower than $10^6$ bacteria/ml. The orange dots correspond to cases with bacterial concentrations larger than $10^6$ bacteria/ml. **a-b** The black lines correspond to the linear regressions applied to low and high concentrations; the variables were logged in base 10. Each linear regression displays the coefficient of determination ($R^2$), the p-value of the slope, and the slope (with standard error).

The change in the power-law exponent (slope in the log-log linear model) for the relationship between phage and bacterial abundances was compared for the communities explored in the model as a function of bacterial density (Figure S.3**a**). The slopes for all communites (same richness) displayed a sublinear exponent ($\alpha < 1$). When compared to the associated bacterial density the value of the slope decreased (linear-log model) with a slope of $\alpha = -0.0334 \pm 0.011$ (p-value$= 0.031 < 0.05$) and a coefficient of determination $R^2 = 0.037$, that is, a weak but statistically significant negative correlation with bacterial density. The change in the relationship between viral and microbial abundances across ecosystems was assessed similarly (Figure S.3**b**). Nine out of the eleven ecosystems displayed a sublinear exponent ($\alpha < 1$). The polar lakes ($\alpha = 1.62 \pm 0.15$) and sediments ($\alpha = -0.07 \pm 0.23$) were the only exceptions. The ecosystems with sublinear exponents displayed a weak but statistically significant negative correlation with microbial density (linear-log regression). The slope was $\beta_1 = -0.17 \pm 0.07$ (p-value$= 0.034 < 0.05$) and the coefficient of determination $R^2 = 0.20$. Thus, individual ecosystems (environmental) and communities (lytic-lysogenic model) displayed sublinear viral-microbial relationships with slopes (power-function exponents) that became flatter upon increase of microbial densities. For most ecosystems and communities, the slopes are smaller than the slop obtained when analyzing data across ecosystems (Figure S.1).

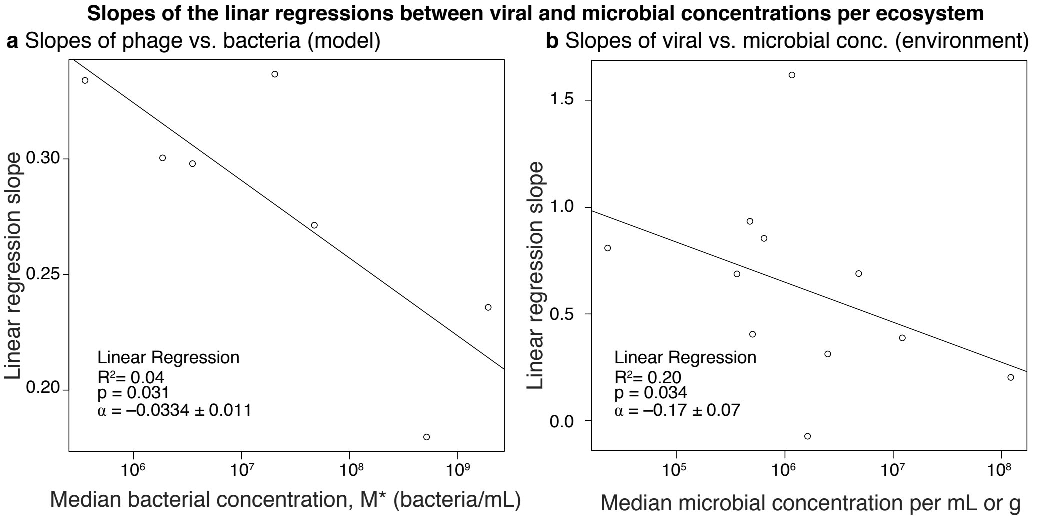


Figure S.3: **Dependence of the power law exponent upon microbial concentration. a** The power law exponents (slopes) obtained in the regressions for communities for fixed richness is plotted as a function of the median bacterial concentration. **a** The power law exponents (slopes) obtained in the regressions for each ecosystem is plotted as a function of the median microbial concentration. **a-b** The black line corresponds to a linear regression. The legend displays the coefficient of determination ($R^2$), the p-value of the slope , and the slope, $\alpha$ (with standard error). In **b** the two outliers with slopes larger than one or smaller than one were excluded from the linear regression.

Table S.1: The file *dataset_ viral_ microbial_ abundances.cvs* contains the reference article, microbial concentration, viral concentration calculated virus-to-microbe ratio, and ecosystem for the environmental data.

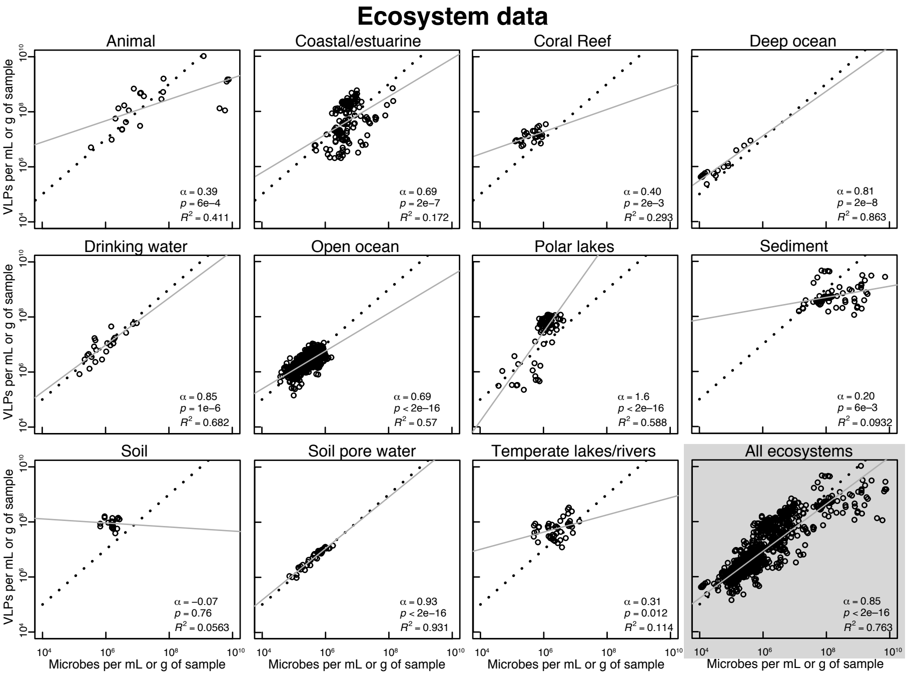


Figure S.4: **Ecosystem Data:** Concentration of viral-like-particles (VLPs) as a function of the microbial concentration combining data from similar ecosystems. The grey line corresponds to the linear regression for the logged data; the power-relationship ($\alpha$), p-value, and R-squared value ($R^2$) are shown in the legend. The dotted line provides a visual reference for the linear case ($\alpha = 1$).

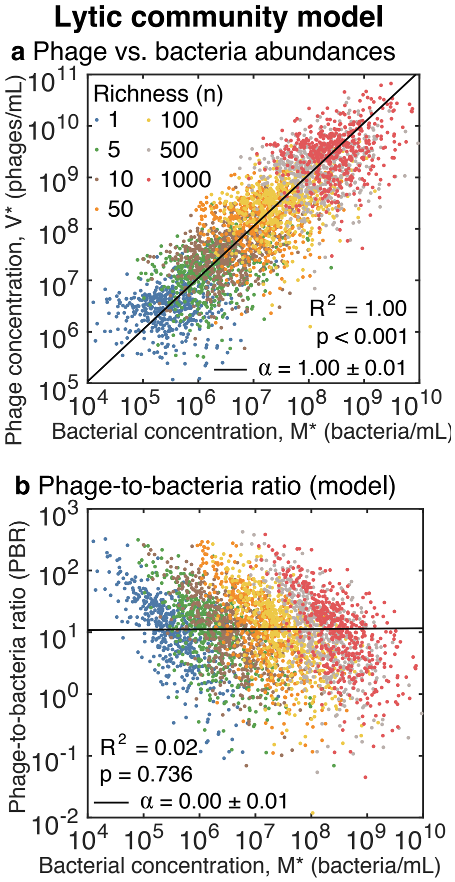


Figure S.5: **Viral and microbial concentrations in the lytic community model.** **a**,Phage equilibrium concentrations versus bacterial concentrations in the ecological lytic community model. The colors distinguish communities with different richness, that is, number of bacteria species or phage-bacteria host pairs. **d**, Phage-to-bacteria (PBR) versus bacterial concentration obtained in the model (same color coding as in panel **a**). **a-b.** The solid lines correspond to the linear regression of the logged data. The slope ($\alpha \pm$ SE), p-value ($\alpha \neq 0$), and the coefficient of determination ($R^2$) are displayed.

#### S.3 EFFECTIVE SINGLE SPECIES MODEL

$$
\begin{aligned}
\frac{\mathrm{d}M}{\mathrm{d}t} &= \overbrace{r\left(1-\frac{M}{K}\right)M}^{\text{logistic growth}} - \overbrace{d'MV}^{\text{infection}}, \\
\frac{\mathrm{d}V}{\mathrm{d}t} &= \overbrace{c'd'MV}^{\text{lysis}} - \overbrace{mV}^{\text{decay}}.
\end{aligned}
\qquad . \tag{S.17}
$$

#### S.4 Sensitivity analysis

A sensitivity analysis was performed by analyzing the relative change of the logarithmic value of the variables with respect each parameter in the model,

$$S_{ij} = \frac{\partial \log y_i^\star}{\partial \log \theta_j}|_\theta = \frac{\theta_j}{y_i^\star}\frac{\partial y_i}{\partial \theta_j}|_\theta \ . \tag{S.18}$$

Here $y_i^\star$ is associated to the equilibrium value of the variables, that is, the total microbial concentration ($y_1^\star = M^\star$) and total viral concentration ($y_2^\star = P^\star$), while $\theta_j$ is associated to the parameters. This are ordered as follows: intrinsic growth rate ($\theta_1 = r$), infection rate constant ($\theta_2 = d$), decay rate constant ($\theta_3 = m$), burst size ($\theta_4 = c$), induction rate constant ($\theta_5 = \beta$), superinfection exclusion factor ($\theta_6 = \chi$), richness ($\theta_7 = n$), and carrying capacity ($\theta_8 = K$). In vectorial form, this reads, $\vec{\theta} = (r, d, m, c, \beta, \chi, n, K)$. The sensitivity factors, $S_{ij}$, were calculated for the mean values of the parameters associated to bacterial and virial physiological traits, a mid-range richness, $n = 100$ and carrying capacity, $K \approx 8,3 \cdot 10^9$. This was associated to approximately 50% of the microbial community being in the lytic compartment and lysogenic compartment. It is worth noting that sensitivity factors are unitless due to the logarithmic approach, which allows the comparison between parameters. The results are independent on the logarithmic based used. The magnitude of the sensitivity factors were ranked for the microbial ($S_{M,j}$) and viral ($S_{V,j}$) communities (Figure S.4). Richness ($n$) was the most relevant parameter for the microbial community ($S_{M,n} \approx 1.5$), followed by the infection rate ($d$), decay rate ($m$), and burst size ($c$), all with sensitivity factors around unity $\approx 1$, while carrying capacity had a sensitivity of half unit ($S_{M,K} \approx 0.5$). The sensitivity of the induction rate ($\beta$), growth rate ($r$), and superinfection exclusion fraction ($\chi$) were negligible in the bacterial community. In the viral community, growth rate ($r$), infection rate ($d$), and richness ($n$) had a similar sensitivity impact around unity ($\approx 1$), while the rest of the parameters were negligible. Thus, richness ($n$) and infection rate ($d$) were sensitive in both the microbial and viral communities, while the other sensitive parameters where phage-trait related in the microbial community (decay rate and burst size) and bacteria-trait related in the viral community (growth rate).

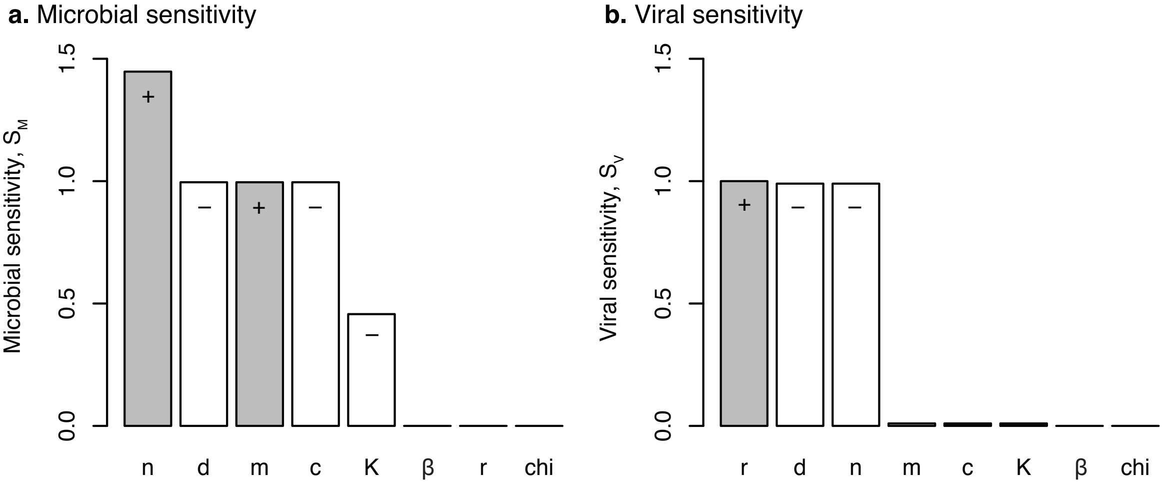


Figure S.6: **Sensitivity analysis.** Response to parameter variations in the microbial community (**a**) and viral community (**b**). The parameters are ranked based on the magnitude of their sensitivity response. Positively correlated responses are displayed in grey (+), while negatively correlated responses are in white (−).
